# Single cell RNA sequencing of *D. pseudoobscura* testes reveals transcriptional signatures of heteromorphic spermatogenesis

**DOI:** 10.64898/2026.07.21.739797

**Authors:** Fiona Messer, April Talbot, Sabrina Williams, Nathan Harmston, Helen White-Cooper

## Abstract

Sperm heteromorphy, the production of multiple sperm morphs with distinct functions, has evolved repeatedly in animals but its developmental and molecular basis remains poorly understood. In the *Drosophila obscura* group, males produce fertilising eusperm and non-fertilising parasperm, yet when and how these lineages diverge is unclear. Here, we applied single-cell RNA sequencing to whole testes of *D. pseudoobscura* to resolve cell-type composition, developmental trajectories, and transcriptional differences between sperm morphs. Analysis of ∼6,500 cells across two replicates identified germline and somatic cell populations and reconstructed spermatogenesis from stem cells to elongated spermatids. We identified a bifurcation in the germline trajectory at the early spermatocyte stage, corresponding to eusperm and parasperm lineages, which was maintained through spermiogenesis. Euspermatocytes, destined to generate longer eusperm, exhibited higher transcriptional activity than paraspermatocytes. Differentially expressed genes included structural sperm tail components, chromatin condensation proteins, post-meiotically transcribed genes and gene duplications resulting in paralogues with reciprocal expression patterns. We also identified markers of somatic cyst cells, including head and tail cyst cells, but found no evidence of morph-specific specialisation within the cyst cell lineage. The transcriptional activities of the distinct germline trajectories corresponded with differential expression of orthologues of critical transcriptional regulators known in *D. melanogaster*. The transcriptional activator *TGIF* was enriched in euspermatocytes, while the transcriptional repressor *kmg* was upregulated in paraspermatocytes. GFP-tagged Kmg showed higher abundance and greater chromatin localisation in paraspermatocytes. These findings demonstrate that sperm morph identity is first apparent in early primary spermatocytes and is largely germline intrinsic. We propose that differential transcriptional provisioning and developmental timing underpin sperm length, providing a framework for understanding the evolution and regulation of sperm heteromorphy.

## Introduction

Sperm heteromorphy is the production of multiple sperm morphs within the same male, with each morph having distinct functions and morphology. Heteromorphic sperm has evolved independently in many phyla, including on multiple occasions in insects, and indeed multiple times in Diptera (Presgraves, et al. 1999; Swallow and Wilkinson 2002; Bhatt 2025).

The *obscura* species group of *Drosophila* all exhibit sperm heteromorphy, producing at least two sperm morphs, eusperm and parasperm, which differ in size (Beatty and Sidhu 1970). Eusperm are longer overall, have longer nuclei, and can fertilise eggs. Parasperm are shorter and not capable of fertilisation (Snook, et al. 1994; Snook and Karr 1998), rather they increase eusperm survival in the female reproductive tract (Holman and Snook 2008; Alpern, et al. 2019). *D. pseudoobscura* produces three sperm morphs: eusperm (∼300μm), parasperm 1 (∼50μm) and parasperm 2 (∼100μm) (Alpern, et al. 2019; Messer and White-Cooper 2025).

Sperm development in the *obscura* species group broadly similar to that of *D. melanogaster* (Lindsley and Tokuyasu 1980; Fuller 1993; Messer, et al. 2026). Sperm production is maintained by the testis niche, comprising the hub, germline stem cells (GSCs) and somatic cyst stem cells (CySCs). The hub acts as a signalling centre that maintains the GSC and CySC populations in an undifferentiated state. GSCs divide asymmetrically; the daughter GSC is retained at the hub, while the daughter gonialblast is displaced from the hub. The gonialblast is encapsulated by two cyst cells, derived from CySCs, forming the spermatogonial cyst, and remains encapsulated for the remainder of spermatogenesis. Cyst cells regulate the cyst’s internal environment, and support germline development, analogous to the function of mammalian Sertoli cells.

Spermatogonia undergo five mitotic divisions with incomplete cytokinesis, resulting in 32 interconnected spermatogonia., which then transition to the primary spermatocyte stage (Kurokawa and Hihara 1976). The primary spermatocytes progress through a rapid S-phase then enter an extended G2 phase, characterised by extensive cell growth and transcription; mRNAs for most of the proteins required after meiosis are transcribed during the spermatocyte stage then stored in a translationally repressed state in the cytoplasm (Olivieri and Olivieri 1965; Schafer, et al. 1995; Barreau, Benson, Gudmannsdottir, et al. 2008; Barreau, Benson and White-Cooper 2008). Primary spermatocytes then undergo the first meiotic division, resulting in a short-lived secondary spermatocyte cyst, which rapidly undergoes meiosis II, to produce the 128-cell spermatid cyst. The spermatid stage is characterised by elongation, nuclear re-shaping, repackaging of the chromatin, individualisation and coiling (Fabian and Brill 2012). The cyst cells differentiate alongside the germline; after meiosis the tail cyst cell ensheaths the elongating tails, while the head cyst cell covers the sperm heads and anchors late spermatid bundles in the testis terminal epithelium. A small number of genes are transcriptionally re-activated in post-meiotic spermatids (Barreau, Benson, Gudmannsdottir, et al. 2008; Barreau, Benson and White-Cooper 2008; Raz, et al. 2023). Many of these have a distinctive localisation to the growing tail-end, termed ‘comet’ and ‘cup’ localisation. Mature spermatozoa are released through the testicular duct into the seminal vesicle at the base of the testis.

All cells within a cyst differentiate into the same morph, suggesting that cell fate is either decided in the gonialblast before the transit amplification, or that a slightly later fate decision is co-ordinately taken by the syncytial germline cells within the cyst (Beatty and Sidhu 1970; Policansky and Ellison 1970; Moore, et al. 2013). Morphological differences between the morphs have not been reported until the spermatid stage, where the spermatids elongate and then begin individualisation.

The genetic underpinning of heteromorphic spermatogenesis is still poorly understood. It is not clear when differentiation between eusperm and parasperm morphs occurs, or what the mechanisms involved in this differentiation process might be. In principle, mechanisms which could act exclusively, or in concert, to determine differential development and result in distinct sperm morphologies include provisioning, timing, blocking and measuring. In a provisioning mechanism, different morphs are provided with different concentrations, compositions or localisations of mRNAs for sperm proteins. Longer eusperm develop from spermatocytes with more RNA provisions, or ones that inherently construct different structures. In a timing mechanism, spermatids elongate for a set time, or morphs have different development rate. For a blocking mechanism, spermatids elongate until they hit a physical barrier, such as the testis wall or terminal epithelium, which issues a ‘stop’ signal. Finally, morphs could have an inherent measuring mechanism, for example a protein gradient, and stop elongating once they reach a set length.

Previous sequencing of single cysts identified differential gene expression between eusperm and parasperm spermatocytes, prior to observing distinct morphological differences between morphs (Messer 2022). This indicates a role for transcriptional control in the development of sperm morphs in *D. pseudoobscura* and supports a ‘provisioning’ hypothesis of heteromorphic sperm development. However, this cyst sequencing provided a limited snapshot of sperm development, and it was unclear from the data which cyst population was destined to develop into which morph. It was also unclear whether cyst cells associated with each morph had distinct identities, whether post-meiotically expressed genes are differentially expressed between the morphs, and what role these factors may have in determining sperm length.

Recently, the FlyCellAtlas and subsequent analysis have demonstrated the huge potential of single-nucleus and single-cell sequencing in the fly testis (Li, et al. 2022; Raz, et al. 2023). Using both nucleus and whole cell sequencing data, Raz, et al. (2023) characterised the developmental trajectories of both germline and cyst cell lineages in the testis, as well as other somatic cell types. Crucially, they were able to map the development of the germline from early proliferating cells through to elongating spermatids, profiling RNA dynamics through each developmental stage. Furthermore, they were able to use the dataset to identify new markers of germline cell types, which were then validated by RNA FISH. They were also able to identify head- and tail-cyst cells based on differential gene expression, validated *in vivo*.

By performing single-cell sequencing on the testis of *D. pseudoobscura*, we have identified and characterised the developmental trajectories of eusperm and parasperm. We demonstrate that morph-enriched differential gene expression is pervasive in the germline from the early spermatocyte stage and identify multiple candidate genes, both transcriptional regulators and sperm structural components, which may have a key role in morph-specific development. Finally, we describe the somatic cell lineages in *D. pseudoobscura* testes, finding no evidence of distinct morph-associated cyst cell populations.

## Methods

### *D. pseudoobscura* strains and culture

Single-cell sequencing was performed on *D. pseudoobscura* wild-type, collected from Show Low, Arizona (SLoB3, gift from Dr. Tom Price, University of Liverpool). To reduce background signal from pigment, we used *white* mutants for HCR-FISH validation (14011-0121.12, National Drosophila Species Stock Centre, Cornell University). Flies were maintained on a standard maize, dextrose, yeast, agar medium at 21°C.

### Single cell sequencing

#### Sample preparation: Testis dissection and cell dissociation

All dissection and dissociation steps were carried out at room temperature. Testes were dissected from *D. pseudoobscura* SLoB3 pupae, at day 7 APF, immediately prior to eclosion. These late pupal testes are attached to the seminal vesicles, have all spermatogenesis stages present (biased towards early stages), and have developed their characteristic bright orange pigmentation.

Testes were dissected on an RNase-free petri dish into a drop of testis buffer (180mM KCl, 40mM NaCl, 10mM Tris pH6.8, in ddH2O), the seminal vesicles, ejaculatory bulb and accessory glands removed, then pooled into a second drop of testis buffer. Dissected testes were washed to remove contaminating tissue, such as fat body, by transferring to fresh 50µl drops of testes buffer, twice. The testes were then transferred to a 50µl drop of dissociation buffer (2mg/ml collagenase, 0.2mg/ml trypsin, DNase I 1unit/ml in testis buffer) on a new petri dish. The testis sheaths were cut open with a sharpened tungsten needle to empty the contents into the dissociation buffer. The pooled sample was mixed with the needle then gently pipetted five times to manually disrupt the sample. The sample was removed to a 1.5ml tube, which had been blocked with BSA. The remaining cells were washed from the dissection plate with a further 50µl drop of dissociation buffer and added to the tube. To manually disrupt the cells, the contents of the tube were gently pipetted for 5 minutes then vortexed on the lowest speed for 1 minute, then pipetted for a further 5 minutes and vortexed for 1 minute. The samples were incubated at room temperature for 10 minutes, pipetted for two minutes, then centrifuged in a swinging bucket rotor at 300g for 5 minutes. The manual disruption steps were repeated a second time. The sample was pipetted for a further 5 minutes, then centrifuged for 5 minutes. The supernatant was removed, and cells resuspended in 100µl dissociation buffer. 5µl of each sample was removed and cell count was estimated using a haemocytometer.

#### Parse Evercode WT: Whole cell fixation, barcoding and library preparation

Cell fixation was carried out immediately after dissociation using the Parse Evercode™ Cell Fixation Kit v2, following the manufacturer’s instructions. For steps requiring a cell strainer, a 70µm mesh size was used, ensuring all stages and cell types were retained in the sample. After fixation, a second 5µl of each sample was removed and cell count estimated using a haemocytometer.

Two replicate samples were prepared, fixed and stored at -80°C prior to barcoding and library preparation. Replicate 1 contained approximately 300,000 cells in a 100μl sample pre-fixation, and approximately 37,500 cells post-fixation, from 41 testes. Replicate 2 contained approximately 508,000 cells in 100μl pre-fixation, and approximately 37,500 cells post-fixation, from 93 testes. Cell barcoding, lysis, cDNA amplification, and library preparation were performed using the Parse Evercode WT Mini kit (v2), according to the manufacturer’s instructions. Barcoded cells were pooled then separated into two ‘sublibrary’ samples. Each sublibrary sample contained cells from both replicates. Subsequent cell lysis and library preparation steps were carried out on both sublibrary samples in parallel.

#### Sequencing and Bioinformatics

Libraries were sequenced by Illumina NovaSeq X (Novogene) using paired end 150bp reads, to a depth of 500 million reads per sample (nominally 50,000 reads per cell).

The raw data were analysed using the Parse Biosciences Splitpipe (v1.2.1) pipeline. QC was performed with FastQC (v0.11.9) (Andrews 2010). Reads were aligned to the *D. pseudoobscura* MV25 reference genome and annotation (GenBank assembly: GCF_009870125.1, Biosample ID: SAMN13616452) with STAR (v2.7.6a) (Dobin, et al. 2013). Reads counts were performed with Samtools (v1.17) (Li, et al. 2009). Data from the two sublibraries were integrated with Splitpipe.

All subsequent bioinformatic and statistical analyses were completed in R Studio (version 4.5.1) (R Core Team 2025), count data were analysed with Seurat (v5.0.1) (Hao, et al. 2024). To remove background sequences, cells were filtered to have a minimum of 399 transcripts per cell. Cells with a transcript count over 250,000 were assumed to be doublets and removed from the dataset. Cells with mitochondrial transcripts over 5% of the total transcript count were assumed to be dying and removed. Counts were log normalised. UMAP dimension reduction was used to plot the data. Germline and somatic cells were separated based on expression of known markers; *always early (aly)* and *polycystine-related-Y (PRY)* for germline, and *eyes absent (eya)* for somatic cell types. Louvain (Louv) cluster analysis of separated germline and somatic data was performed at resolutions 0.8, 1.6 and 3.2. Differential gene expression analysis between clusters was performed using the Wilcoxon Rank Sum test with FindAllMarkers in Seurat. Pairwise cluster comparisons were performed using the FindMarkers function.

#### Motif analysis

To identify enrichment of transcription factor binding motifs in eusperm and parasperm-enriched genes, we performed motif analysis of differentially expressed genes between eu- and paraspermatocytes, at early (clusters 20 and 21, Louv 3.2) and mid-spermatocyte stages (clusters 7 and 9, Louv 0.8). Promoters were defined +/- 150bp centred on the annotated TSS. Motif enrichment analysis was performed using monaLisa for *D. melanogaster* motifs present in the JASPAR database, using a binomial test and a minimum score of 80% (Machlab, et al. 2022; Rauluseviciute, et al. 2024). Two additional motifs were also included in this set of motifs, representing the CNAAATT and Aly motif (Katzenberger, et al. 2012; Kim, et al. 2017; Lu, et al. 2020). Motifs with an FDR < 0.1 were defined as significantly enriched in either eusperm or parasperm.

#### HCR fluorescent *in situ* hybridisation

We validated our manual annotations of the scRNAseq dataset by HCR-FISH of whole adult testes (Choi, et al. 2018; Moth, et al. 2024; Shao, et al. 2024). Probes were designed using the ‘probe picker’ tool (Matthew Jachimowicz, University of Toronto). A full list of probes can be found in supplementary methods.

Testes were dissected into testis buffer then fixed in 0.5% PFA in PBT (PBS + 0.1% Tween 20) for 10 minutes, followed by a further 30 minutes fixation in 4% PFA in PBT. Testes were then washed twice with 1ml PBT, for 5 minutes. Fixed testes were pooled and stored in methanol at -20°C for a minimum of 16 hours.

The methanol was removed and the samples rinsed with PBT, followed by 100μl 30% probe hybridisation buffer (30% formamide, 5X sodium chloride sodium citrate (SSC), 9mM citric acid (pH 6.0), 0.1% Tween 20, 50μg/ml heparin, 5X Denhart’s solution, 10% dextran sulfate). Testes were pre-hybridised for 30 minutes at 37°C in 100μl 30% probe hybridisation buffer. Probe solutions were prepared by adding 0.4μl of 100μM probes to 100μl of 30% probe hybridisation buffer. Both probe solution and pre-hybridised testes in probe hybridisation buffer were transferred to a heat block set to 80°C for 5 minutes. The pre-hybridisation buffer was removed from the testes and replaced with the probe solution. The testes in probe solution were incubated for a further 5 minutes at 80°C, then transferred to a 37°C heat block overnight.

Probe solution was replaced with 30% probe wash buffer (30% formamide, 5X SSC, 9mM citric acid (pH 6.0), 0.1% Tween 20, 50μg/ml heparin). Testes were washed a total of 5 times, 20 minutes per wash, at 37°C. Wash buffer was removed and replaced with amplification buffer (5X SSC, 0.1% Tween 20, 10% dextran sulfate). Testes were incubated in amplification buffer for 30 minutes at room temperature. 2μl per sample of each hairpin was heated at 95°C for 90 seconds, then left to cool at room temperature for 30 minutes. Hairpin solutions were prepared by adding 2μl of each heated and cooled hairpin to 50μl amplification buffer. Testes were incubated in hairpin solution overnight at room temperature, in the dark.

Testes were rinsed once, then washed 4x 5 minutes with 5X SSCT (5X SSC, 0.1% Tween 20). They were washed 2x 5 minutes, with PBS, and a final 15 minute was with PBS + Hoechst 33258, to visualise the DNA. Testes were transferred to mounting media on slides (70% glycerol, 2.5% n-propyl gallate, 0.1% Hoechst 33258). HCR-FISH samples were imaged with a Zeiss LSM880 confocal microscope. Image analysis was performed using ImageJ.

#### Transgenic *D. pseudoobscura* expressing a Kmg-GFP fusion protein

We constructed a piggyBac vector containing a selectable AmCyan eye marker, and the sequence for C-terminally tagged Kmg-GFP, for random insertion at a TTAA site in the genome (Handler, et al. 1998; Holtzman, et al. 2010; Navarro Paya 2017).

Kmg fragments were amplified from synthesised template (IDT). We included 1000bp upstream of the TSS to ensure we captured the Kmg promotor. Primers for the Kmg promotor, 5’UTR and CDS fragment included extensions for the AvrII restriction site, primers for the 3’UTR included NotI extensions. GFP sequence was amplified from *D. melanogaster w;UAS-eGFP-TEVt-RFP-M4/CyO* (Jianqiao Jiang) with a 5’ AvrII restriction site extension. Amplified fragments were ligated into pGEM-T Easy. GFP and *kmg* fragments were ligated sequentially into a piggyBac-3xP3-AmCyan backbone using AvrII and NotI sites (Navarro Paya 2017) (supplementary methods). Insert orientation was confirmed by sequencing (Eurofins).

An injection mix of 500ng/μl piggyBac-Kmg-GFP:300ng/μl piggyBac helper was injected into *D. pseudoobscura* SLoB3 embryos. *F_0_* survivors were collected and crossed to *D. pseudoobscura* SLoB3. *F_1_*flies were screened for AmCyan eyes/ocelli and crossed with *D. pseudoobscura* SLoB3. Transgenic *F_2_* flies were backcrossed, and homozygous *F_3_* collected to set up stable stocks. *D. pseudoobscura* Kmg-GFP testes were imaged by Zeiss LSM880 confocal microscopy. Image analysis was performed using ImageJ.

### Sequence analysis of *D. pseudoobscura* prominin-like gene family

Paralogues of *prominin-like (promL)* in *D. pseudoobscura* were identified by PSI-BLAST of LOC4812066, the syntenic orthologue of *D. melanogaster promL* (Altschul, et al. 1997; Schaffer, et al. 2001). Coding sequences for 13 *D. pseudoobscura*, two *D. melanogaster* and one *Aedes aegypti* promL proteins were retrieved from NCBI and aligned with Clustal-Omega (Madeira, et al. 2024). Phylogenetic trees were generated with FastTree (2.1.11) (Price, et al. 2010).

## Results

We performed single cell RNA sequencing of whole *D. pseudoobscura* testes dissected from 7-day old pupae, across two replicates. We aimed to characterise *D. pseudoobscura* testis transcription, with an emphasis on characterising the developmental trajectories of eusperm and parasperm sperm morphs and identifying the molecular mechanisms underpinning differential morph development. For clarity, where we have referred to genes by the name of their *D. melanogaster* orthologue, the *D. pseudoobscura* LOC gene number can be found in Supplementary Table 1.

### Overview: Whole testis data contains both germline and somatic cell types

After filtering, a total of 6534 cells were analysed across two replicates (Rep 1 = 2714, Rep 2 = 3820), with 16880 features detected (fig. 1A). Median transcript count per cell across all data was 4273 (min = 461, max = 245012), median feature count per cell was 1790.5 (min = 399, max = 9713) (fig 1B-C). Clustering of the whole dataset at Louvain resolution 0.8 resulted in 15 clusters which were separated into germline and somatic cell types for subsequent analysis by expression patterns of four known markers (Li, et al. 2022; Raz, et al. 2023) (fig 1D, fig S1A-B). Germline clusters were indicated by *always early (aly)* and *polycystine-related-Y (PRY)* and somatic clusters by *Myosin heavy chain (Mhc)* and *eyes absent (eya)* (fig 1D-H). The data were separated based on germline vs. somatic classification for all subsequent analysis (fig 1I-N). Louvain clustering was performed separately on both germline and somatic datasets, at three resolutions, 0.8, 1.6 and 3.2 (fig 1I-N). Germline clusters 0 and 1 (Louvain 0.8) both showed low transcript and feature counts (median < 1000) and were excluded from subsequent analysis, as ‘cells’ in these clusters are likely to be ambient RNA artefacts or debris from the library preparation (Supplementary Table 2). Both replicates were represented in all clusters of germline and somatic data (Supplementary Table 2, 3).

**Figure 1:**
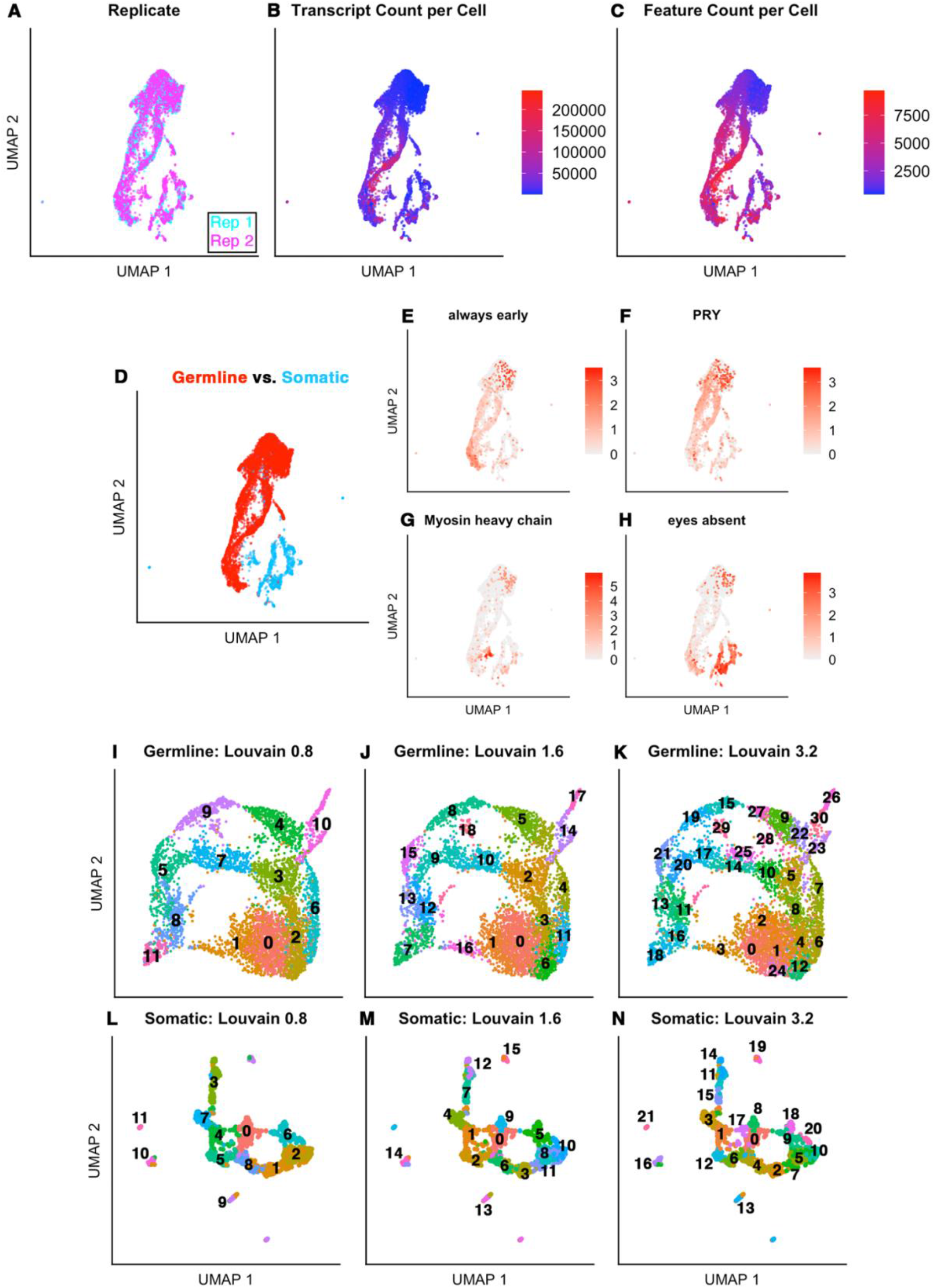
Overview of *D. pseudoobscura* whole testis single cell RNA sequencing data. Dimension reduction by UMAP with Louvain clustering. (A) Both replicates show equal distribution across the UMAP, indicating replicability of the data. (B-C) Transcript and feature count per cell. (D) UMAP of the whole testis dataset showing germline and somatic cell types. (E-H) UMAPs showing expression of germline and somatic cell markers. (E-F) Early germline marker *always early*, and late germline marker *polycystine-related-Y*, showing higher expression in the larger collection of clusters (1, 7-15) indicating germline cells. (G) *Myosin heavy chain* indicating muscle cells in cluster 2, and (H) *eyes absent* indicating cyst cells in clusters 3-6. (I-N) Germline and somatic cells were separated and dimension reduction and clustering algorithms performed independently on the two datasets. (J-L) Germline UMAP with Louvain clustering at three resolutions; 0.8, 1.6 and 3.2. (M-O) Somatic UMAP with clustering at Louvain 0.8, 1.6 and 3.2 resolutions.

### Germline cell types: Clustering identifies germline developmental stages

To assign identities to clusters, we first examined broad expression patterns of known marker genes in relation to lower resolution clustering, at Louvain 0.8. Higher resolutions (1.6 and 3.2) showed broadly the same patterns of clustering, but with further subdivisions into smaller clusters (fig 1J-L, fig 2A-B, fig S1, Supplementary Table 4). As expression patterns of markers are likely to differ in *D. pseudoobscura*, compared to those characterised in *D. melanogaster*, we also validated cluster identity by RNA *in-situ* by HCR-FISH of whole mount testes.

Cluster identities are summarised in figure 2 (A, B) and Table 1. Markers for each cluster, based on those used in the FlyCellAtlas testis dataset, are provided as a dot matrix in figure 2C and are described in detail below (Li, et al. 2022; Raz, et al. 2023).

**Figure 2:**
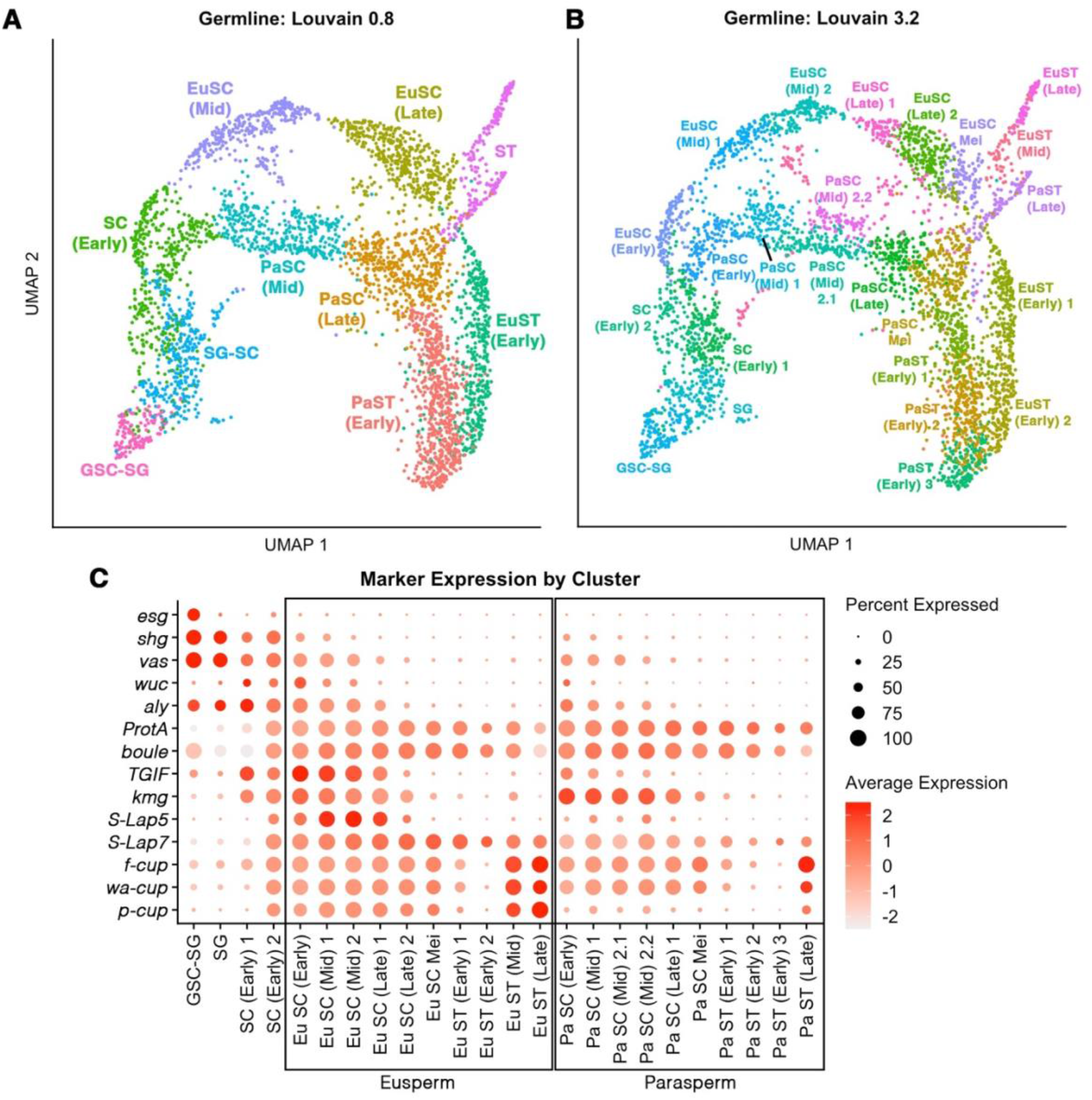
Summary of germline data. (A-B) UMAP reduction and Louvain clustering was applied to cells assigned germline identity. Louvain clustering resolutions at 0.8 and 3.2. Higher resolution resulting in greater subdivision of clusters. Cluster identity was assigned based on expression of known cell-type markers in *D. melanogaster* and validated by HCR-FISH. (C) Expression of markers by cluster at Louvain resolution 3.2, ordered by developmental stage and morph. Eusperm and parasperm data are separated after the trajectory split at the early spermatocyte stage.

**Table 1:**
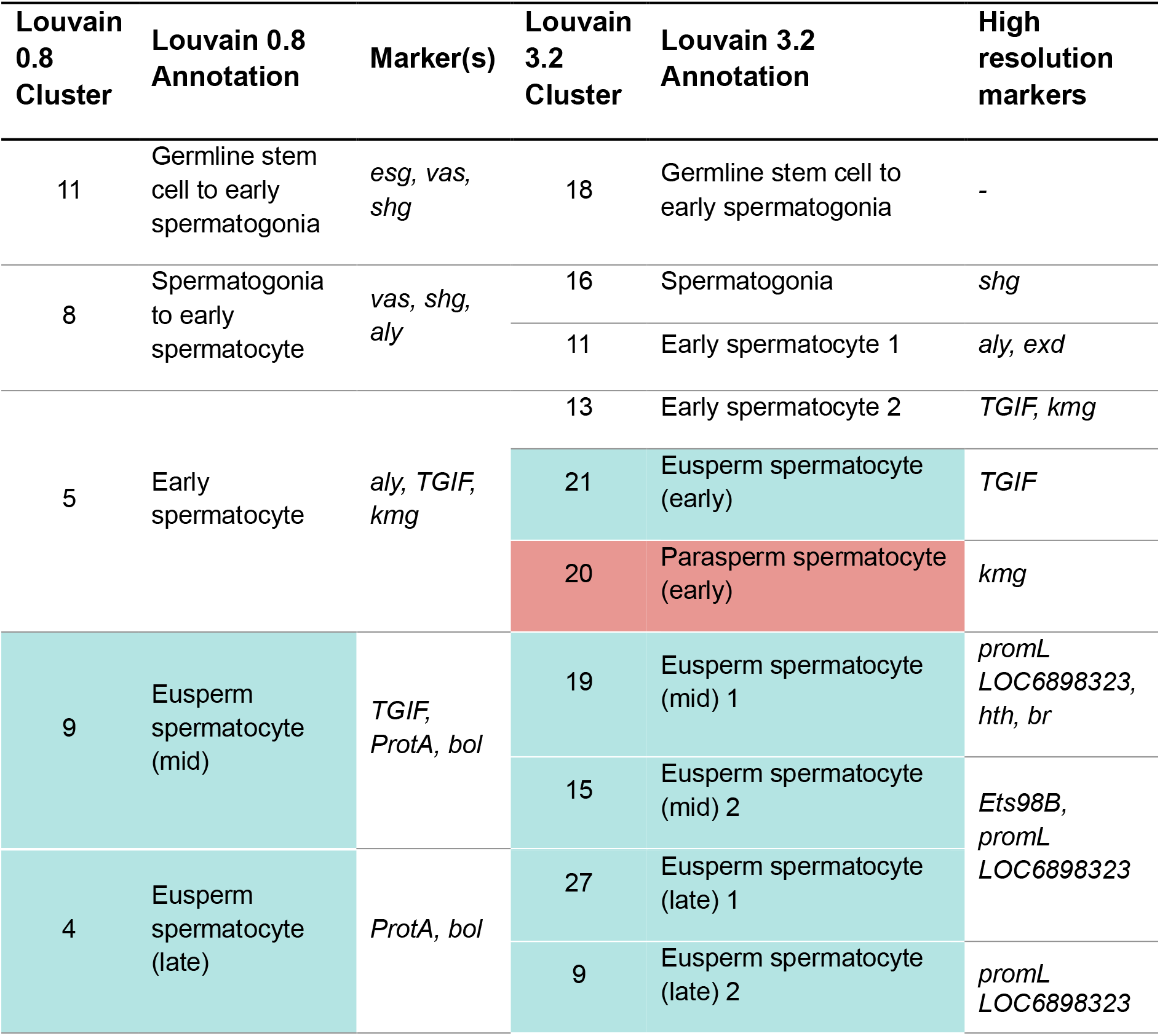
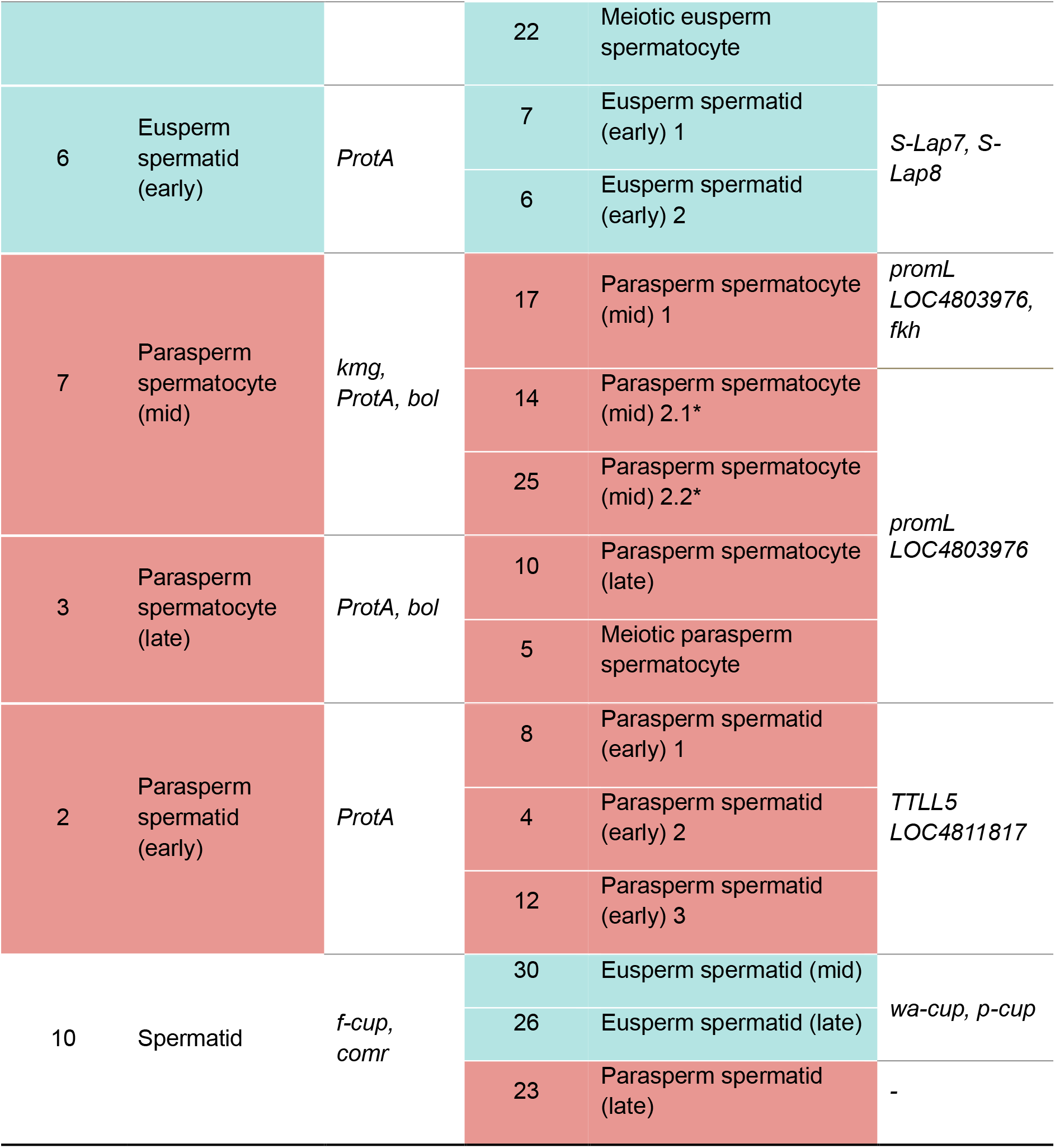
Germline cluster annotations at Louvain resolutions 0.8 and 3.2. As clustering was broadly stable, higher resolution clusters are indicated as subdivisions of lower resolution clusters. Clusters assigned to the eusperm trajectory are indicated in blue, clusters assigned to the parasperm trajectory are indicated in red. Marker genes for Louvain 0.8 cluster annotations are indicated. Additional markers and differentially expressed genes in higher resolution Louvain 3.2 clusters are indicated. *A second trajectory split in the mid paraspermatocyte stage may represent parasperm 1 and parasperm 2 morphs.

### Germline stem cells, spermatogonia and early spermatocytes

*escargot (esg)* is a marker for germline and somatic stem cells, proliferating spermatogonia, and the hub. In the germline data, *esg* expression was limited to cluster 11 (Louvain 0.8), indicating that these are germline stem cells and proliferating spermatogonia (fig 3A).

**Figure 3:**
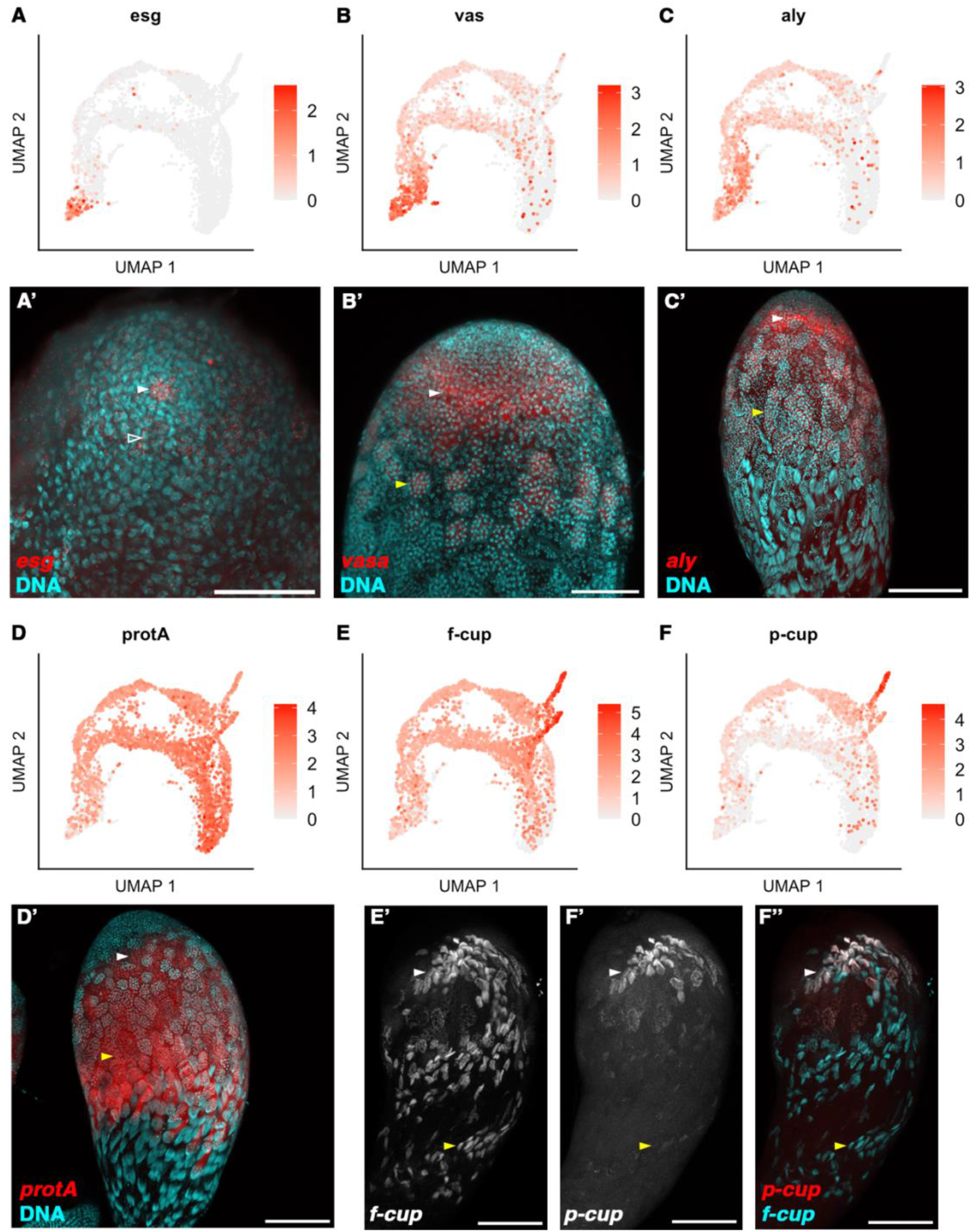
Validation of germline cluster annotation. (A-F) UMAPs of germline stage marker genes. (A’-F’’) HCR-FISH staining showing RNA localisation in whole mount *D. pseudoobscura w^-^* testes. (A, A’) Hub, germline stem cell (GSC) and spermatogonia marker *escargot*. Arrowhead indicates hub. Open arrowhead indicates spermatogonia. (B, B’) GSC and spermatogonia marker *vasa*. (C-C’) Primary spermatocyte marker *always early*. (D, D’) Late primary spermatocyte and early spermatid marker *protamine A*. (E, E’) Elongating spermatid marker *flyers cup*. (F, F’) Late elongating euspermatid marker *presidents-cup*. (F’’) Double HCR-FISH of *f-cup* and *p-cup*, showing eusperm-specific transcription and localisation of *p-cup*. (B’-D’) White arrowhead indicates eusperm primary spermatocyte cyst. Yellow arrowhead indicates parasperm primary spermatocyte cyst. (E’-F’’) White arrowhead indicates euspermatid tail, yellow arrowhead indicates paraspermatid tail.

*vasa (vas)* is also marker of GSCs and spermatogonia and shows expression in clusters 11 and 8 (Louv 0.8) (fig 3B). HCR-FISH confirmed that *vas* is maintained later than *esg*, into the early spermatocyte stage, indicating cluster 8 may contain later spermatogonia (16 cell) transitioning to the early spermatocyte stage (32-cell). *vas* decreases in clusters 5, 7 and 9, suggesting that the these are later germline stages.

Cluster 8 (Louv 0.8) also shows an increase in expression of *aly* (fig 3C), a key component of the testis meiotic arrest complex (tMAC) required for testis-specific gene expression in primary spermatocytes. *D. melanogaster aly* is expression peaks in early primary spermatocyte and maintained until late spermatocyte stage (White-Cooper, et al. 2000; Raz, et al. 2023).

At a higher resolution cluster 8 (Louv 0.8) is subdivided into two clusters, 16 and 11 (Louv 3.2) (fig 1J, K, Table 1). A dot matrix shows cluster 16 maintains higher expression of stem cell and spermatogonia markers *shotgun (shg) and vas*, with early spermatocyte markers *aly, TGIF* (orthologous to *D. melanogaster achi* and *vis*) and *kumgang (kmg)* increasing in cluster 11, and further spermatocyte genes expressed in cluster 13. We therefore suggest the cluster 8 (Louv 0.8) contains both spermatogonia and early spermatocytes, the spermatogonia to spermatocyte transition occurring between clusters 11 and 13 (Louv 3.2).

### Germline differentiation splits into two trajectories at the spermatocyte stage

Cluster 5 (Louv 0.8) shows both high expression of early spermatocyte markers (*aly, TGIF, kmg*), and increasing expression of later spermatocyte genes, including *protamine A (protA)* and *boule (bol*). In *D. melanogaster, protamine A (protA)* is transcribed in the mid-primary spermatocyte stage, later than *vas* and *aly,* and it depends on *aly* for expression (Raz, et al. 2023). Clusters 5, 7 and 9 (Louvain 0.8) show increasing expression of *protA*, peaking in clusters 3, 4, 2, 6 and 10 (Louvain 0.8) (fig 3D). HCR-FISH also shows high levels of *protA* increasing from the early spermatocyte stage, peaking as the spermatocytes enter meiosis and maintained through secondary spermatocytes, early spermatids, until mid-elongation. We conclude that clusters 5, 7 and 9 comprise early to mid-spermatocytes, while clusters 3 and 4 are later spermatocytes.

At higher resolution, cluster 5 (Louv 0.8) separated into three clusters, 13, 20, and 21 (Louv 3.2) (fig 1J, K). Cluster 13 lies earliest in the differentiation trajectory, splitting to generate clusters 20 and 21. Spermatocyte genes *TGIF, kmg, S-Lap5* and *S-Lap7* were differentially expressed after this split. The split is maintained through two major trajectories, the ‘upper’ trajectory (Louv 3.2 clusters 21, 19, 15, 27, 9), and a ‘lower’ trajectory (Louv 3.2 clusters 20, 17, 14, 25, 10). The lower trajectory has a secondary split, between clusters 14 and 25, although this split is not maintained into later stages.

### Post-meiotic transcription in elongating spermatids confirms eusperm and parasperm trajectory identities

Post-meiotic stages are typically characterised by overall low levels of active transcription, although RNA transcribed during the spermatocyte stage is present in spermatids, stored in RNPs until translation. A subset of genes transcribed post-meiosis show low levels of transcription during the spermatocyte stage, ceasing as the primary spermatocytes enter meiosis, followed by a marked increase in transcription in spermatids. Many of these genes also show distinctive localisation patterns to the spermatid distal tail-end, termed ‘cup’ or ‘comet’ localisation (Barreau, Benson, Gudmannsdottir, et al. 2008; Barreau, Benson and White-Cooper 2008; Raz, et al. 2023).

*flyers cup (f-cup), walker cup (wa-cup) and president’s cup (p-cup)*, are post-meiotically transcribed in *D. melanogaster*, and their mRNAs localise to the distal tail-ends of spermatids (Barreau, Benson, Gudmannsdottir, et al. 2008; Barreau, Benson and White-Cooper 2008). Cluster 10 (Louvain 0.8) showed high levels of *f-cup*, indicating that these are elongating spermatids (fig 3E). Clusters 2 and 6 showed comparatively low levels of *f-cup* indicating that these are early spermatids.

Clustering at Louvain 3.2 separated the elongating spermatids into three clusters, 30, 26 and 23. Clusters 30 and 26 are connected to the ‘upper’ spermatocyte trajectory, whereas cluster 23 is connected to the ‘lower’ spermatocyte trajectory. Clusters 30 and 26 show high levels of post-meiotic genes *f-cup, wa-cup* and *p-cup*, while cluster 23 had high *f-cup,* but comparatively low expression of *wa-cup* and *p-cup*. HCR-FISH showed *f-cup* was present in both eu- and paraspermatids, whose identity can be inferred from their position in the testis. Spermatid heads are oriented towards the base of the testis and their tails elongate internally within the testis towards the apical region. The 300µm tails of the long euspermatids are long enough to reach close to the testis tip, while those of the shorter paraspematids are found more basally in the testis. *p-cup* was high in euspermatids, but low or not detectable in paraspermatids (fig 2F). This indicates that the upper trajectory corresponds to eusperm development, and the lower trajectory to parasperm development. We did not identify two parasperm trajectories maintained through to the elongation stages, however the secondary split in the lower trajectory (Louv 3.2 clusters 14 and 25) may be indicative parasperm 1 and 2 clusters.

### Differential gene expression between eusperm and parasperm trajectories

Having identified that the trajectory split between eusperm and parasperm occurs at the early spermatocyte stage, we further characterised differential gene expression between comparable developmental stages. Both transcript count and feature count were higher in the eusperm trajectory, increasing from cluster 13 (Louv 3.2, SC Early 2) and continuing as the trajectory splits, decreasing by the meiotic stage (cluster 22, Louv 3.2). Transcript count and feature count in the parasperm trajectory paralleled this pattern, but both were lower overall than the eusperm trajectory (fig 4A-B, Supplementary Table 2-3). Many genes upregulated in euspermatocytes initiate transcription in cluster 13 (Louv 3.2, SC Early 2), potentially pinpointing the earliest split between eusperm and parasperm trajectories.

**Figure 4:**
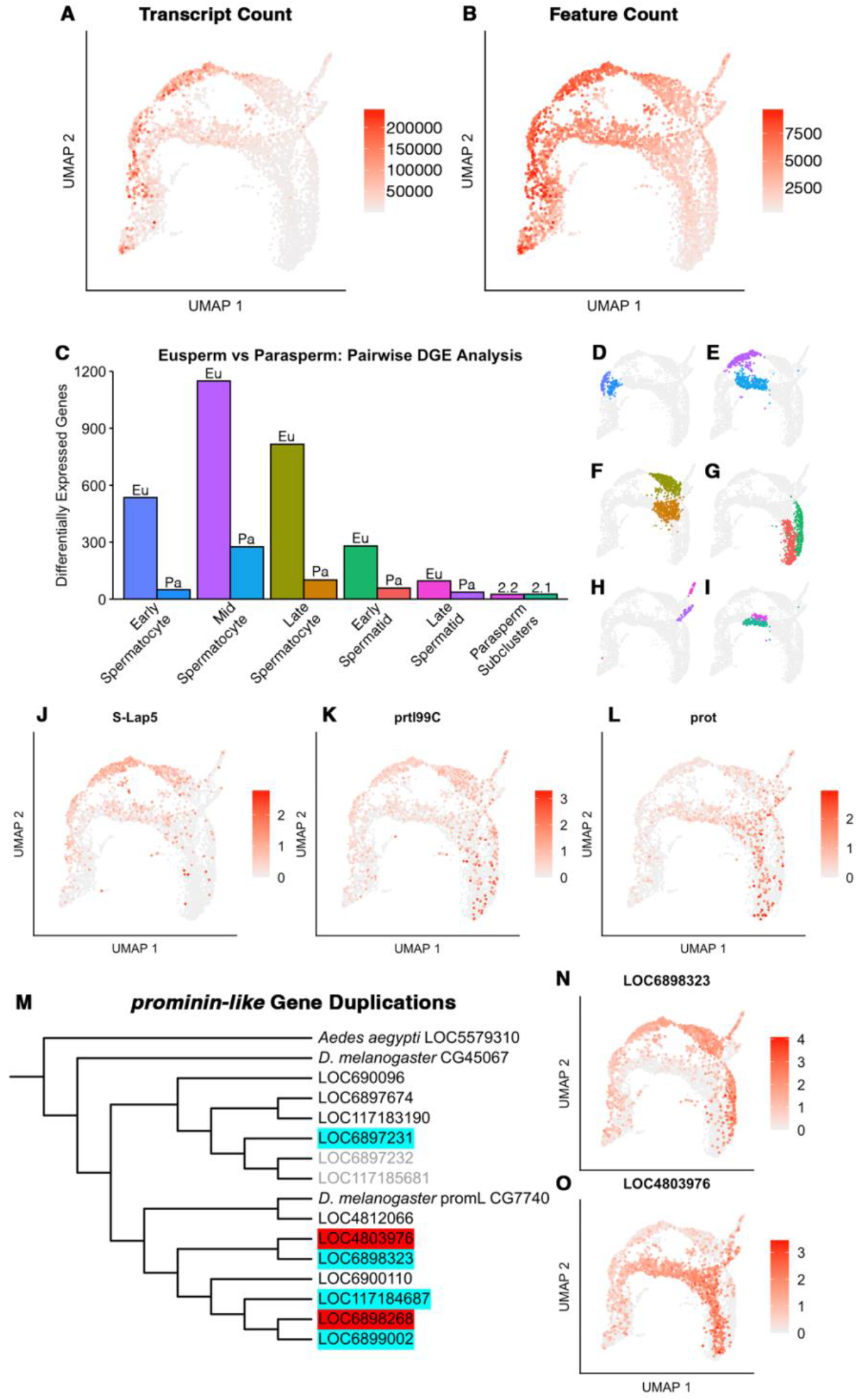
Differential gene expression analysis of germline scRNAseq. (A-B) UMAPs of transcript count and feature count. (C-I) Summary of differential gene expression analysis between eusperm and parasperm clusters of equivalent developmental stage. Across all developmental stages, a greater number of differentially expressed genes were upregulated in eusperm clusters. Parasperm subclusters had a negligible difference in upregulation of DE genes. (D-I) UMAPs of clusters compared for differential gene expression analysis. (D) Early spermatocyte, (E) mid-spermatocyte, (F) late-spermatocyte, (G) early spermatid, (H) late spermatid, (I) parasperm sub-clusters. (J) Upregulation of *Sperm-Leucylaminopeptidase 5* in eusperm spermatocytes. (K) Upregulation of *protamine-like SSC* in euspermatocytes. (L) Upregulation of *protamine* in paraspermatocytes. (M-O) Duplication and differential expression of *prominin-like (promL)* genes in *D. pseudoobscura*. (M) Maximum-likelihood cladogram tree of 13 *D. pseudoobscura promL* genes shows many duplication events have occurred in *D. pseudoobscura*, while a single duplication is present in *D. melanogaster*. Cyan indicates upregulation in eusperm trajectory, red indicates upregulation in parasperm trajectory. Genes in light grey were not expressed in either the germline or somatic cells of the testis. (N-O) UMAPs a *promL* tandem pair, *LOCc8S8323* and *LOC4803S7c*, showing upregulation in eusperm and parasperm trajectories respectively.

We performed pairwise differential gene expression analysis, directly comparing clusters of equivalent developmental stage between the eusperm and parasperm trajectories. We found a greater number of differentially expressed genes were upregulated in the eusperm trajectory, compared to the parasperm trajectory, at all stages (fig 4C-I, Supplementary Table 5). In early euspermatocytes, 535 genes were upregulated (greater than 2-fold change, adjusted p-value <0.05), compared to 50 in early paraspermatocytes. This rose to 1149 in mid-euspermatocytes, compared to 275 in mid-paraspermatocytes, decreasing in late euspermatocytes to 816, compared to 101 in late paraspermatocytes. The trend continued in spermatid stages; 280 genes upregulated in early euspermatids, compared to 58 in early paraspermatids, and 96 genes in late euspermatids, compared to 37 in late paraspermatids.

We also noted that paraspermatocyte cysts are positioned lower in the testis than eusperm cysts of the equivalent stage, suggesting that they develop at a slower rate.

#### Sperm tail components show differential gene expression between eusperm and parasperm spermatocytes

Given that largest morphological distinction between eusperm and parasperm morphs is their tail length, it may be expected that genes encoding tail components be differentially expressed, with upregulation in the eusperm. Consistent with this, we identified three sperm leucyl aminopeptidases (S-Laps) upregulated in euspermatocytes, *S-Lap5, S-Lap7* and *S-Lap8* (fig 4J, fig S2). S-Laps are structural components of the paracrystalline material in the mitochondrial derivatives; these structures extend the length of the tail and are thus more extensive in eusperm. Also higher in eusperm were *CG302c8* (*LOC480384c*), predicted in *D. melanogaster* to be a structural component of the radial spokes (Zur Lage, et al. 2019), and cilium assembly/intraflagellar transport components *BBIP1* and *rempA* (Lee, et al. 2008). The canonical intraflagellar transport mechanism is required for differentiation of sensory cilia, but not spermatogenesis (Han, et al. 2003), however BBIP1 and RempA may have alternate functions in spermatogenesis, unrelated to intraflagellar transport, or *D. pseudoobscura* may require a level of intraflagellar transport for assembly of the eusperm tail.

Conversely, we identified a smaller number of genes with potential tail-assembly function upregulated in parasperm; two *TTLL5* paralogues and *Pericentrin-like protein (Plp)* (Lee, et al. 2008). The function of TTLL5 in spermatogenesis is not known, although other tubulin tyrosine ligases function in post-translational modification of tubulins, thereby modulating stability. In *D. melanogaster*, *Plp* maintains the structure of the pericentriolar material, thereby contributing to axoneme structure (Martinez-Campos, et al. 2004; Richens, et al. 2015). Interestingly, several core sperm tail components did not show evidence of differential expression. The axonemal β2-tubulin gene, *βTub85D*, showed high expression across both morphs, but was not significantly differentially expressed (fig S2). Of the ancestrally Y-linked kinesin-like genes, required for sperm motility, *kl-2* showed no evidence of differential gene expression, although both *kl-3* and *kl-5* were marginally upregulated in euspermatocytes (Log2FC = ∼0.5, p<0.001). Given that these are the key structural components of the sperm tail, it perhaps surprising that these were not observed to be more differentially expressed. Instead, there may be some post-transcriptional mechanisms which differentially regulate the translation of these RNAs during the elongation process, or their expression is not rate-limiting for growth.

#### Some sperm head components appear to have morph enrichment

Unlike sperm tail components, we expected that genes required for DNA condensation during spermiogenesis would not show differential gene expression, as both eusperm and parasperm contain the same quantity of DNA, although their morphology is distinct (Pasini, et al. 1996). Indeed, a major component of the condensed chromatin, *protamine A*, showed no differential gene expression (fig 3D). However, two other protamines did show significant differential expression; *protamine-like SSC* was upregulated in eusperm, and *protamine* was enriched in parasperm, suggesting that there is some specialisation of essential head components between morphs (fig 4K, L).

#### Gene duplication and subfunctionalisation between eusperm and parasperm

A striking example of morph enrichment of paralogues was that of the prominin-like genes. The function of prominins in sperm development has not been well characterised, although there is evidence that they are required for sperm development in mammals, and may localise to the membrane at the ends of developing sperm tails (Fargeas, et al. 2004; Matsukuma, et al. 2023). In *D. melanogaster*, Prominin-like is known to have functions in multiple somatic tissues and functions: maintaining the architecture of highly curved membranes of extracellular vesicles and microvilli, eye development, insulin signalling, mitochondrial function, and locomotion (Mahato, et al. 2018; Ryu, et al. 2019; Wang, et al. 2019; Zheng, et al. 2019; Hurbain, et al. 2022; Ryu, et al. 2022).

Two prominin-like proteins are present in *D. melanogaster*, *prominin-like* (*PromL*) and *CG45067*. In *D. pseudoobscura* we identified 13 prominin-like proteins, of which six were differentially expressed between morphs (fig 4M-O, fig S3). Four were upregulated in euspermatocytes, with two upregulated in paraspermatocytes. Evolutionary distance did not appear to influence which morph the prominin-like was specialised to; in two instances, very recently duplicated pairs were upregulated in different morphs.

#### Post-meiotic gene expression

While most gene expression in spermatogenesis occurs in the pre-meiotic spermatocytes, a subgroup of genes are (re)activated post-meiosis, in the spermatids. 162 such genes have been identified in *D. melanogaster* (Barreau, Benson, Gudmannsdottir, et al. 2008; Barreau, Benson and White-Cooper 2008; Raz, et al. 2023). To identify post-meiotic expressed genes in *D. pseudoobscura*, performed differential gene expression analysis between late spermatids (cluster 10, Louv 0.8) and early spermatids (combined clusters 2 and 6, Louv 0.8). We identified 203 genes upregulated in late spermatids; orthologues of 34 of these had also been identified as post-meiotically expressed in *D. melanogaster.* Of the 203 *D. pseudoobscura* post-meiotically expressed genes, 63 were upregulated in euspermatids, whereas none were upregulated in paraspermatids (Supplementary Table 5).

#### Parasperm 1 and 2 show limited transcriptional differences

Pairwise comparison between the two putative paraspermatocyte morph clusters (clusters 14 and 25, Louv 3.2, Figure 4I, Table 1) revealed low differential gene expression; 51 genes with greater than 2-fold change and adjusted p-value <0.05. These were evenly split between the two clusters; 25 upregulated in paraspermatocyte 2.1 (cluster 14) and 26 in paraspermatocyte 2.2 (cluster 25). If these clusters are indeed representative of the two parasperm morphs, this indicates that parasperm 1 and 2 are transcriptionally very similar during the spermatocyte stage of development, when the highest levels of transcription occur. This is also reflective of similar morphology between parasperm 1 and 2 (Alpern, et al. 2019; Messer and White-Cooper 2025).

### Transcriptional upregulation in eusperm spermatocytes may be regulated by testis transcription factor complexes tMAC and tTFIID

Euspermatocytes show higher transcription and a greater number of genes expressed. To investigate the underlying mechanism, we investigated known testis transcription factor genes for evidence of differential expression between eu- and paraspermatocytes.

#### Transcriptional regulators TGIF and wake-up call are upregulated in euspermatocytes

The *D. melanogaster* testis meiotic arrest complex (tMAC) is a pioneer transcription factor complex required for opening chromatin at the promotors of spermatocyte-expressed genes (Ayyar, et al. 2003; Wang and Mann 2003; Beall, et al. 2007; Laktionov, et al. 2014; Laktionov, et al. 2018; Lu, et al. 2020). Mutants for tMAC components fail to undergo both meiosis and spermatid differentiation (Beall, et al. 2007; Messer, et al. 2026). As tMAC is essential for transcription in spermatogenesis, we validated expression patterns of all tMAC subunits and known binding partners by HCR-FISH (fig 5). *Caf1* is a single copy gene in *D. melanogaster* but has two paralogues in *D. pseudoobscura*. *always early (aly), chromatin assembly factor 1B (caf1B), cookie monster (comr), myb-interacting protein 40 (mip40), TGIF, tombola (tomb), matotopetli (topi)* and *wake-up call (wuc)* were transcriptional activated in early spermatocytes (cluster 11, Louv 3.2). *chromatin assembly factor 1A (caf1A)* showed high expression in spermatogonia, decreasing in early spermatocytes, followed by re-activation in early eu- and paraspermatocytes after the trajectory split (cluster 20 and 21, Louv 3.2). Surprisingly, *comr* also showed post-meiotic expression in both eu- and paraspermatids, localising to the spermatid tail ends (fig 5D). This post-meiotic expression and localisation pattern is not observed in *D. melanogaster comr* (Jiang and White-Cooper 2003; Raz, et al. 2023).

**Figure 5:**
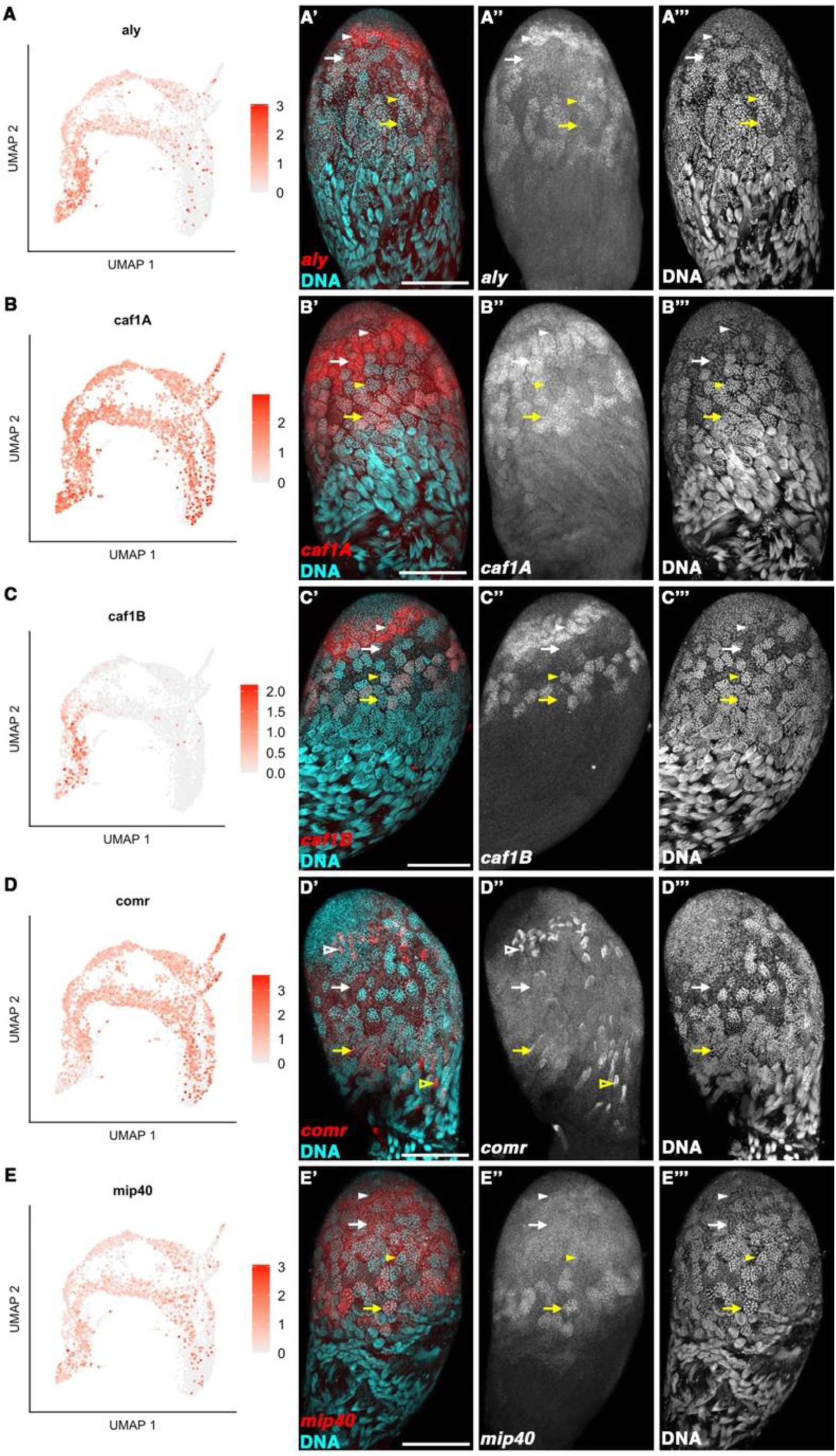
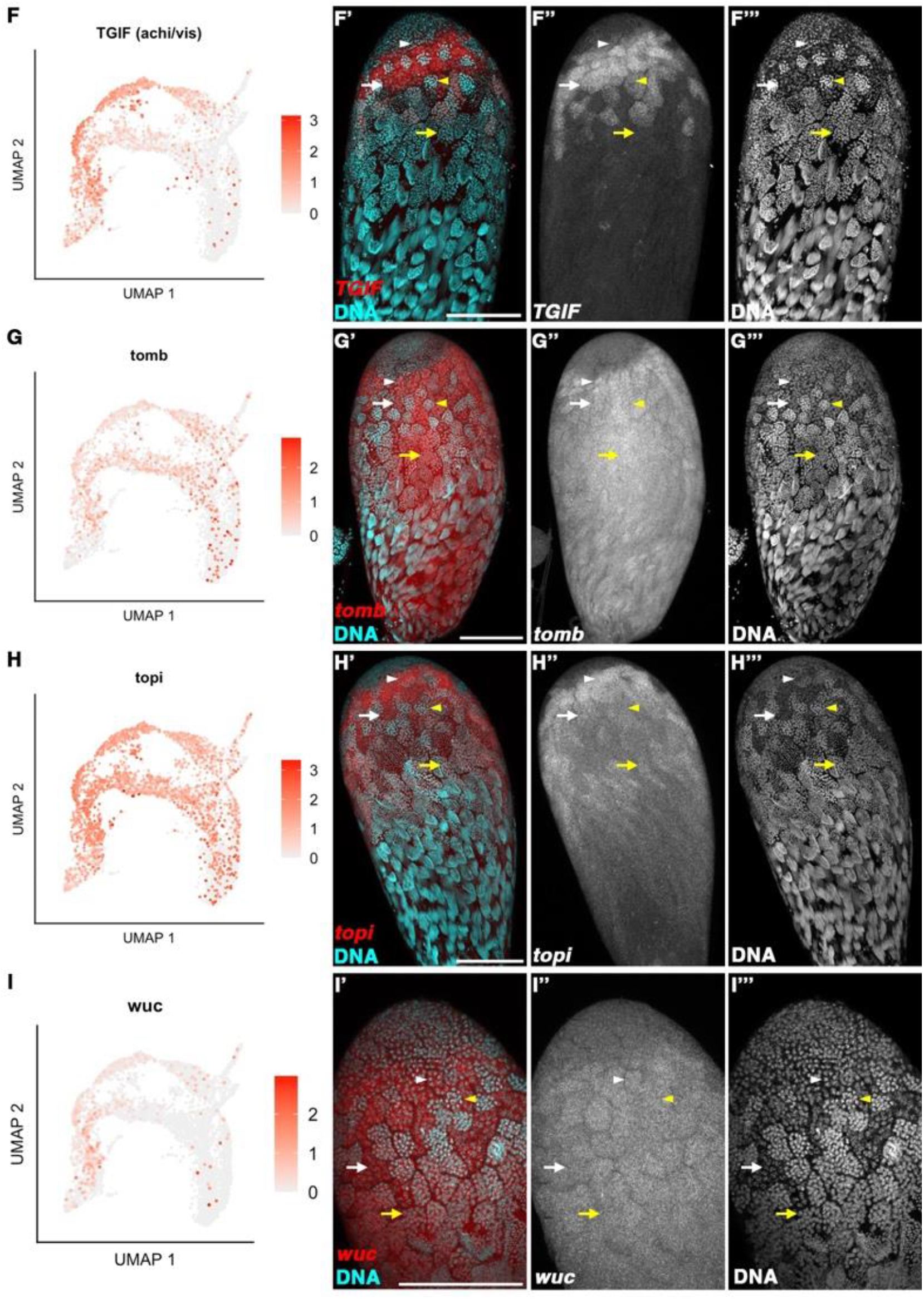
Orthologues of tMAC subunits and associated proteins. (A-I) UMAPs of TMAC and associated proteins. (A’-I’’’) HCR-FISH in whole mount *D. pseudoobscura w^-^* testes. Arrowheads indicate early primary spermatocyte cysts, arrows indicate mid to late primary spermatocyte cysts, open arrowheads indicate spermatid cysts. White indicates eusperm cysts, yellow indicates parasperm cysts. (A-A’’’) *aly* is high in early spermatocytes, then declines in mid-spermatocytes. (B-B’’’) *caf1A* is present throughout germline development, showing upregulation in the early-mid spermatocyte stage. (C-C’’’) *caf1B* is expressed in early spermatocytes, decreasing in mid-spermatocytes. (D-D’’’) *comr* is expressed in pre-meiotically in primary spermatocytes, and in post-meiotic elongating spermatids, localising to the tail-ends of spermatid cysts (open arrowheads). (E-E’’’) *mip40* is expressed in pre-meiotic germline, then decreasing in meiotic and early spermatid cysts. (F-F’’’) *TGIF (achi/vis)* is transcribed in early spermatocyte cysts prior to the trajectory split, and is then upregulated in euspermatocytes, declining in late-euspermatocytes. (G-H’’’) *tomb* and topi are upregulated in early primary spermatocytes and maintained until spermatid elongation. (I-I’’’) *wuc* is marginally upregulated in euspermatocytes, declining in mid-late euspermatocytes.

Two tMAC-associated proteins were significantly upregulated in euspermatocytes; *TGIF* and *wuc*. *TGIF* (orthologues *achi* and *vis D. melanogaster*) was upregulated in early and mid-euspermatocytes. A UMAP plot of *TGIF* showed high expression in early spermatocytes prior to the trajectory split (cluster 13, Louv 3.2), maintaining at a high level in euspermatocytes until meiosis (fig 5F). *TGIF* was also present in paraspermatocytes, but at a lower level. This was confirmed by *in situ* hybridisation; *TGIF* signal was higher in euspermatocytes and was present in late euspermatocytes. Paraspermatocytes had lower signal and was not detectable above background in late paraspermatocytes. *wuc* was upregulated in mid-euspermatocytes, although the transcript was detected in less than 40% of cells at this stage, and differential expression between eu- and paraspermatocyte cysts was not clear in FISH (fig 5I).

#### TGIF is a sequence-specific DNA binding protein that co-immunoprecipitates with core tMAC subunits

tMAC is a testis-specific complex paralogous to the somatic MMB/DREAM complex (Beall, et al. 2007; White-Cooper and Davidson 2011). *Mip130,* the paralogue of *aly* that predominantly functions in soma in *D. melanogaster*, was also upregulated in euspermatocytes (Beall, et al. 2007; White-Cooper and Davidson 2011) (fig S4).

#### TFIID subunits and their testis-TFIID paralogues show differential expression in spermatocytes

The general transcription complex, TFIID comprises TBP and TBP-associated proteins (TAFs); testis specific paralogoues of TAFs (tTAFs) likely generate a testis specific complex, tTFIID(Hiller, et al. 2001; Hiller, et al. 2004; Metcalf and Wassarman 2007). The tTAF *nht* was upregulated in euspermatocytes. *Taf5*, the TFIID paralogue of the tTFIID component *can*, and *Taf7*, another TFIID component, were also upregulated in euspermatocytes (fig S4), as was *tbrd-1*, a testis-specific bromodomain containing protein (Theofel, et al. 2014).

### Kumgang, a transcriptional repressor of tMAC-dependent gene expression, is upregulated in parasperm spermatocytes

While differentially expressed testis transcription factor genes were almost all upregulated in euspermatocytes, we found *kumgang (kmg)*, a transcriptional repressor, was upregulated in paraspermatocytes (Kim, et al. 2017). *kmg* showed an increase in expression from the early spermatocyte stage (cluster 13, Louv 3.2). *kmg* was marginally upregulated in early paraspermatocytes (cluster 20, Louv 3.2, Log2FC = 0.75), and significantly upregulated in mid-paraspermatocytes (cluster 7, Louv 0.8, Log2FC = 1.25). *In situ* hybridisation confirmed this expression pattern and showed *kmg* transcript persisted into early paraspermatids (fig 6A-B). To confirm that *kmg* was upregulated in a different subset of cysts than tMAC transcription factor *TGIF*, we performed a double *in situ* hybridisation. A subset of early spermatocyte cysts showed high levels of *TGIF* with a medium level of *kmg*, while the remaining cysts, positioned more basally, had high levels of *kmg* with low *TGIF*. In the late spermatocyte cysts, a subset of cysts had low expression of both *TGIF* and *kmg*, while the other late spermatocyte cysts had no detectable *TGIF* above background, but *kmg* remained high (fig 6C). This confirms that *TGIF* and *kmg* were upregulated in different morphs.

**Figure 6:**
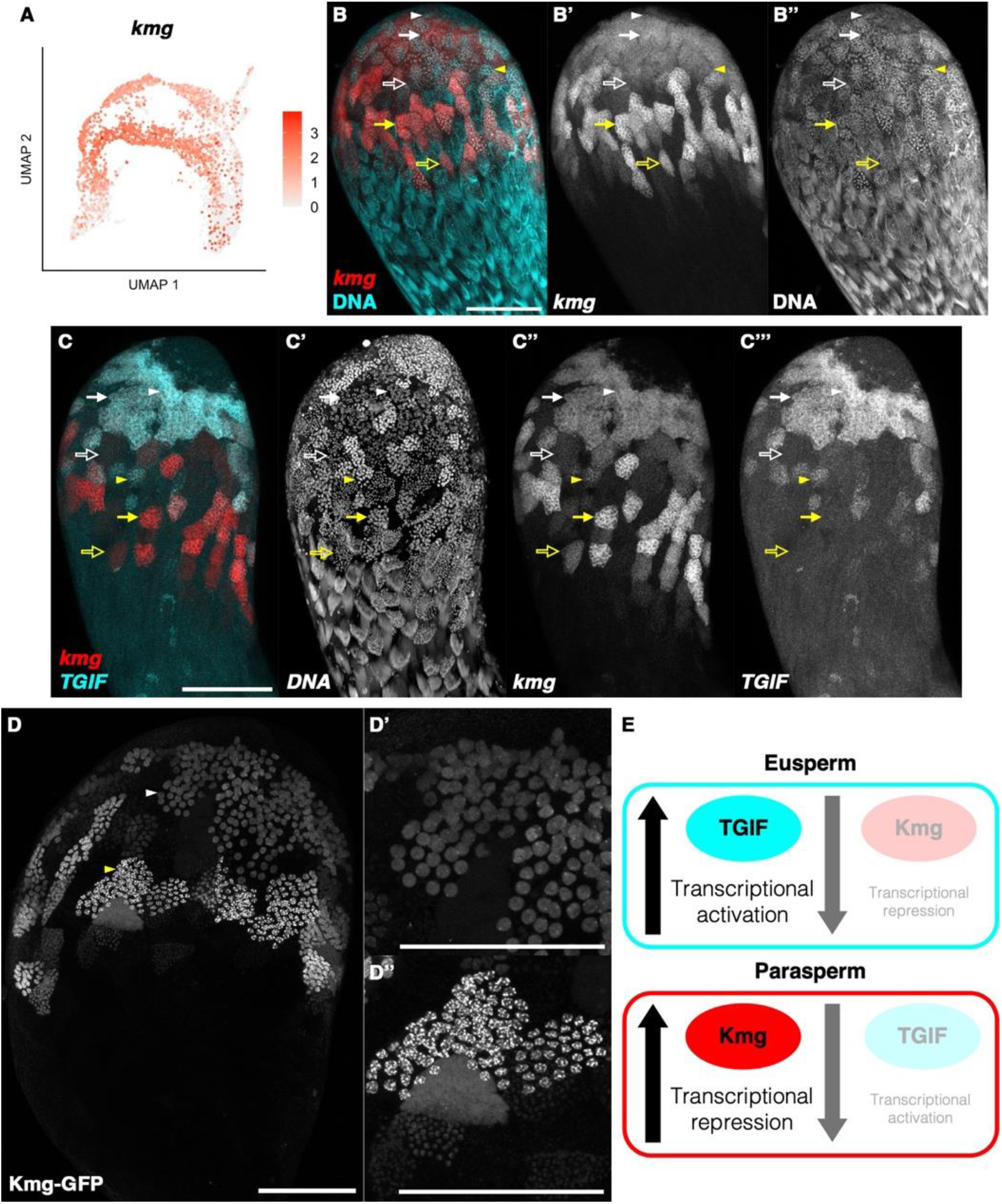
Transcriptional repressor Kumgang (Kmg) expressed in primary eu- and paraspermatocytes but is upregulated in paraspermatocytes. (A) UMAP of *kmg* showing upregulation in early, mid and late paraspermatocytes. (B-B’’) HCR-FISH for *kmg* in whole mount *D. pseudoobscura w^-^* testes. (C-C’’’) HCR-FISH double staining for *kmg* and *TGIF* showing high levels of both *kmg* and *TGIF* in early eusperm and parasperm spermatocyte cysts. TGIF is higher in mid-euspermatocyte cysts compared to mid-paraspermatocyte cysts. Conversely, *kmg* is higher in mid-late paraspermatocyte cysts. (B-C’’’) Arrowheads indicate early primary spermatocyte cysts, arrows indicate mid primary spermatocytes cysts, open arrows indicate late primary spermatocyte cysts. White indicates eusperm cysts, yellow indicates parasperm cysts. (D-D’’) Kmg-GFP protein fusion shows upregulation of Kmg in paraspermatocyte cysts. White arrowhead indicates euspermatocyte cyst, yellow arrowhead indicates paraspermatocyte cyst. (D’) Euspermatocyte cysts. (D’’) Paraspermatocyte cysts show higher levels of Kmg-GFP and greater localisation to chromatin. (E) Models of Kmg and TGIF mechanisms to regulate higher transcription in euspermatocytes and restrict transcription in paraspermatocytes.

*D. melanogaster* Kmg prevents tMAC- (and TGIF) dependent transcription from cryptic promoters of around 2000 genes which are either normally expressed in the soma or not expressed in any tissue, thereby preventing their expression in the germline (Kim, et al. 2017; Matias, et al. 2025).

*D. pseudoobscura* Kmg may have a similar function in early eu- and paraspermatocytes. The precise level of tMAC target gene expression in spermatocytes involves a dynamic battle between tMAC promoting chromatin opening, RNA polymerase recruitment and transcription initiation, and Kmg (with NuRD) counteracting this. This involves interaction between Kmg, NuRD components and the nascent RNA (Matias, et al. 2025). *D. pseudoobscura kmg* was upregulated in paraspermatocytes, indicating it may tip the balance towards lower tMAC target gene expression during parasperm development.

To further explore a potential role for Kmg in paraspermatocyte transcriptional repression, we generated a transgenic *D. pseudoobscura* line expression a Kmg-GFP fusion protein, by *piggyBac*-mediated random insertion. Kmg-GFP was expressed in spermatocytes, with expression resembling that of the RNA (fig 6D). The slightly larger, more apically positioned euspermatocyte cysts showed nuclear localisation of the fusion protein, with 1-2 puncta per nucleus. The slightly smaller, more basal paraspermatocyte cysts also showed nuclear localisation of Kmg-GFP but also had higher overall levels of Kmg-GFP, and at least five bright puncta per nucleus, consistent with chromatin localisation. Kmg-GFP also persisted at a higher level in the parasperm secondary spermatocytes and early spermatids. This supports a role for Kmg in tuning down transcription in paraspermatocytes, while high levels of transcriptional activation in euspermatocytes is mediated through TGIF and tMAC, as in *D. melanogaster* (fig 6E).

*D. melanogaster* Kmg acts with chromatin remodelling factors Mi-2, Dany and Simj (Kim, et al. 2017; Matias, et al. 2025). While all three were expressed in *D. pseudoobscura* spermatocyte clusters, neither *mi-2* nor *simj* were differentially expressed between morphs. Surprisingly, *dany* was marginally upregulated in early and mid-euspermatocytes (Louv 3.2 cluster 21 Log2FC = 1.0, p = 0.15; Louv 0.8 cluster 9 Log2FC = 0.99, p<0.001) (fig S5). *Mi-2*, *dany* and *simj* may have a function in differential morph transcription regulation via their interactions with Kmg, but this is unlikely to be driven by their differential abundance in eu- and paraspermatocytes.

### Motif analysis identified additional regulatory mechanisms

To identify additional potential transcription factor involvement, motif enrichment analysis was conducted of promoter regions of genes enriched in either eusperm or parasperm (fig 7, fig S6, Supplementary Table 6). The TCE motif (CNAAAT), which is enriched in tMAC target promoters in

**Figure 7:**
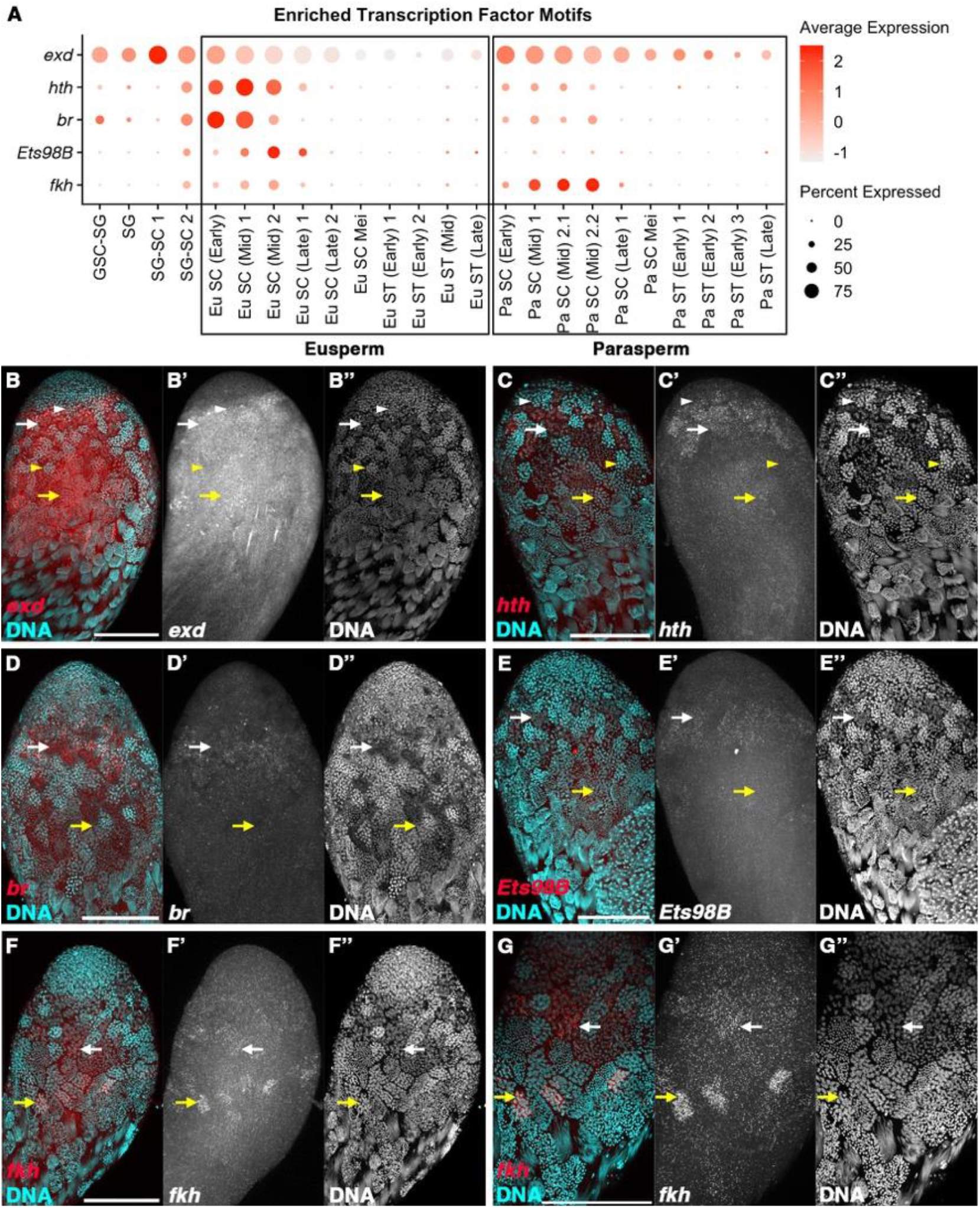
Expression of transcriptional regulators identified by motif analysis of eu- and paraspermatocyte clusters. (A) Dot matrix plot of differentially expressed putative transcription regulators. (B-G’’) HCR-FISH validation of transcriptional regulators. Arrowheads indicate early primary spermatocyte cysts; arrows indicate late primary spermatocyte cysts. White indicates euspermatocyte cysts, yellow indicates paraspermatocyte cysts. (B-B’’) *extradenticle* is marginally upregulated in paraspermatocytes. (C-E’’) *homothorax, broad* and *EtsS8B* were upregulated in euspermatocytes. (F-G’’) *forkhead* was expressed in late primary spermatocyte cysts and upregulated in late-paraspermatocytes compared to late-euspermatocytes.

*D. melanogaster*, was enriched in promoters of early euspermatocyte-biased genes, consistent with a role in the initial divergence of the trajectories (fig S6A). An enrichment for the *TGIF*-motif (*vis)*, was observed in late paraspermatocyte-biased genes (fig S6D).

In eusperm-upregulated genes at early timepoints, promoters were enriched for *homothorax (hth)* and *forkhead (fkh)* motifs, as well as motifs bound by PRD-like homeodomain transcription factors, suggesting coordinated early regulatory control. At later timepoints, eusperm-upregulated genes showed enrichment for E-box motifs, potentially indicating a shift in regulatory inputs during development (fig S6B). Overall, a greater number of enriched motifs were identified in eusperm compared to parasperm, indicating that the regulation of eusperm enriched genes is more complex than parasperm.

Most of the transcription factors binding to these putative regulatory motifs are either not expressed in male germline, or, expressed at a low level but not differentially between the lineages. We performed FISH for those that were identified as differentially expressed between eu- and paraspermatocytes in the scRNAseq (fig 7). *hth, broad (br)* and *Ets98B* transcripts were higher in euspermatocytes, consistent with them potentially acting to upregulate targets in this lineage (fig 7C-E). In contrast, *extradenticle (exd)* and *fkh* motifs were enriched in euspermatocytes, but the genes were themselves upregulated in paraspermatocytes. This would suggest they have a repressive role, lowering expression of euspermatocyte-enriched genes in paraspermatocytes (fig 7A-B, F-G).

### The testis somatic cell populations

In addition to germline, the testis contains multiple somatic cell types, which are essential for germline maintenance and development. As the somatic dataset contained fewer cells than the germline data, we assigned cluster identity based on the lower resolution clustering (Louv. 0.8). Cluster identity assignment of the somatic dataset is summarised in figure 8A.

**Figure 8:**
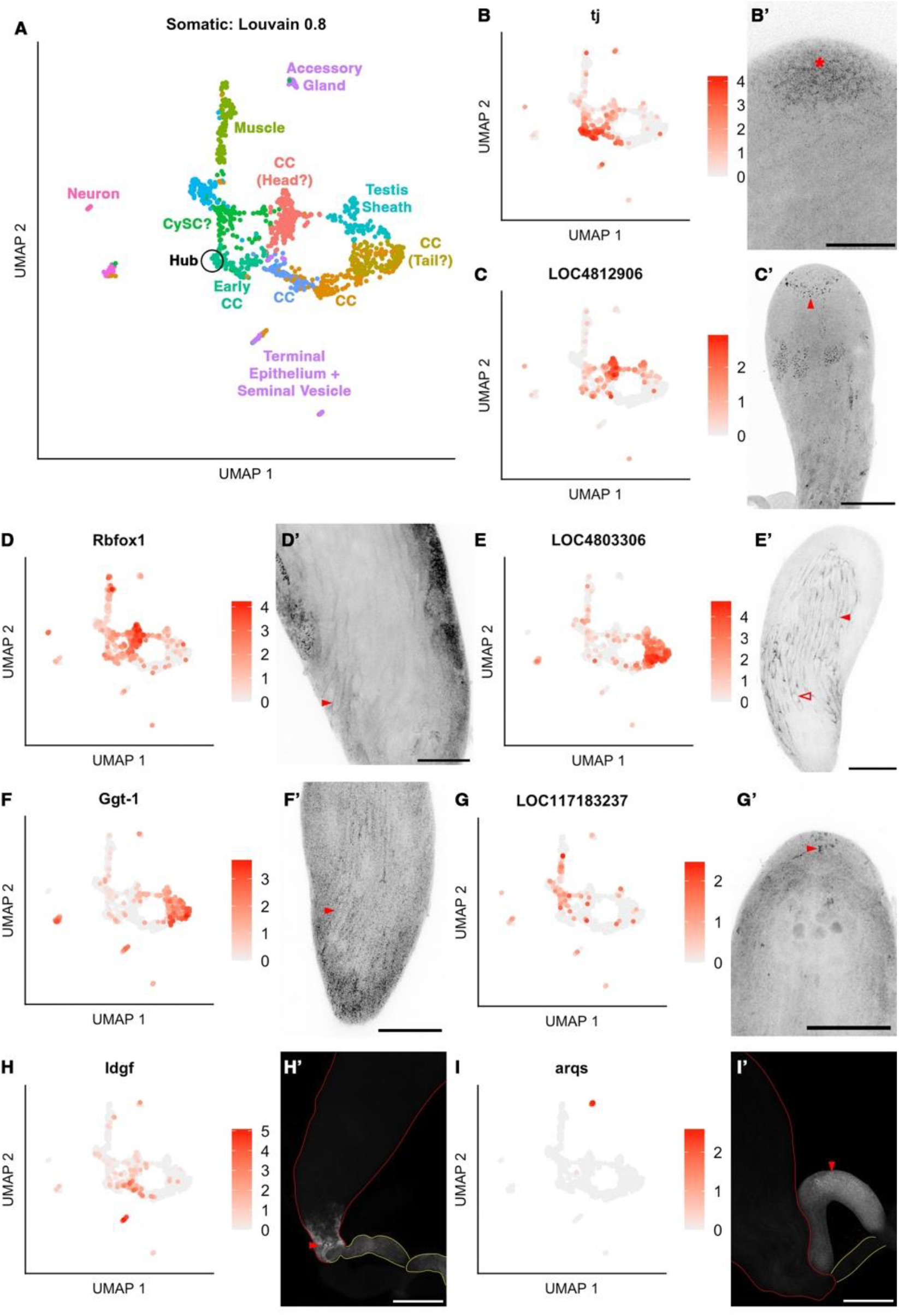
Somatic cell cluster annotations and validation. (A) UMAP of somatic cells, clustered at resolution Louvain 0.8. (B-I) UMAPS of somatic markers genes, either previously identified in *D. melanogaster* or identified from this dataset. (B’-I’) HCR-FISH of marker genes validates somatic cluster annotation. Images are presented using inverted grayscale for clarity. (B-B’) *traffic jam* is a marker of hub, cyst stem cell and early cyst cells. Asterisk indicates hub. (C-C’’) *LOC4812S0c* marks cyst cells around the spermatogonia-spermatocyte transition region of the testis (arrowhead). It is also present in early spermatid tails. (D-D’) *Rbfox1* identified head cyst cells of late spermatid cysts (arrowhead) and also has early spermatid tail localisation. (E-F’) *LOC480330c* and *Ggt-1* identify tail cyst cells of spermatid cysts (arrowheads). Outlines of waste bags can also be identified in E’ (open arrowhead). (G-G’) *LOC117183237* identified cyst cells close to the hub region of the testis and may indicate cyst stem cells (arrowhead). (H-H’) *Idgf* marks terminal epithelium (arrowhead), testicular duct and seminal vesicle. (I-I’) *aqrs* is expressed in accessory gland (arrowhead). (H’-I’) Red outline indicates testis, yellow outline indicates testicular duct and seminal vesicle.

#### Head and tail cyst cells have distinct trajectories

The cyst cell clusters within the somatic data were indicated by previously characterised cyst cell markers *traffic jam (tj), eyes absent (eya)* and *sine oculis (so)* (Fabrizio, et al. 2003; Wingert and DiNardo 2015). However, cyst cells were difficult to visualise with FISH for *eya* and *so*; we identified additional cluster markers using Seurat and validated cluster identity by FISH. Cyst cell trajectories were also informed by 3D UMAP of the somatic data (fig S7).

Early cyst cells were identified by *tj* (Wingert and DiNardo 2015). *tj* was upregulated in cluster 5 (Louv 0.8), and FISH for *tj* showed localisation to cyst cells in most apical region of the testis, consistent with cluster 5 containing cyst stem cells and early cyst cells (fig 8B).

In *D. melanogaster, eya* is upregulated in mid-stage cyst cells, while *so* is upregulated in late cyst cells. Both were upregulated in cluster 0 (Louv 0.8), indicating that this cluster represented later cyst cells than cluster 5 (fig S8). As neither *eya* nor *so* gave clear FISH signal, we identified two genes upregulated in cluster 0 with low expression in other somatic clusters.

LOC4812906 is orthologous to *D. melanogaster* CG32052, a neuronal gene with low expression in the testis, mostly restricted to the germline. FISH showed a ring-like region of signal, near the apical tip of the testis, immediately basal to the region stained by *tj* (fig 8C). Signal was also detected in early spermatids, although this was not observed in the germline scRNAseq dataset. A second marker, *Rbfox1*, was also upregulated in cluster 0. Like LOC4812906, FISH signal was observed in early spermatids but was also localised to the head cyst cells of elongating spermatid cysts (fig 8D). Cluster 0 may therefore contain head cyst cells ranging in developmental stage, associated with spermatocytes through to elongating spermatids.

We were able to identify tail cyst cells with markers for cluster 2 (Louv 0.8). *LOC4803306* (orthologous to *CG312Sc*) and *Ggt-1* were both expressed in tail cyst cells of elongating spermatids (fig 8E, F). *CG312Sc* is upregulated in mid-cyst cells in *D. melanogaster* but has no known function. Ggt-1 is an enzyme involved in glutamate and aspartate biosynthesis, which also shows late cyst cell expression in *D. melanogaster* (Li, et al. 2022; Raz, et al. 2023).

Clusters 8 and 1, which connect clusters 5 (early cyst cells) and 2 (elongating tail cyst cells), showed lower transcript and feature counts than the other clusters in the somatic dataset. We were unable to identify markers which distinguished between clusters 8 and 1, or from other cyst cell clusters. These clusters may represent earlier tail cyst cells than cluster 2. Alternatively, these may represent broken cyst cells; broken cells would be expected to show an overall similar transcriptional pattern, with lower transcript counts compared to intact cells of the same type. Cyst cells associated with elongating spermatids are extremely long and thin, increasing the likelihood of cell breakage during cell dissociation and fixation.

*LOC117183237* is a non-coding RNA upregulated in cluster 4 (Louv 0.8). FISH signal was localised to the apical tip region of the testis, indicating that these may be cyst stem cells (fig 8G). *tj* is also a marker of cyst stem cells and showed a low level of expression in cluster 4, as did *LOC4812S0c*. *LOC117183237* and *LOC4812S0c* had a more punctate expression than *tj*, which was more diffuse, but all were localised to the apical region of the testis.

While we were able to identify head and tail cyst cell clusters, we did not identify morph-specific cyst cell populations. Validation of the cyst cell lineages by FISH did not identify any markers of eusperm or parasperm-associated cyst cells, suggesting that cyst cells of eusperm and parasperm cysts are not transcriptionally distinct.

#### Seminal vesicle and accessory glands

While seminal vesicles and accessory glands were removed from the testes during dissection, it is likely that a small number of cells from these tissues were present in the dissociated cell suspensions. Cluster 9 (Louv 0.8) comprises three smaller clusters, which at higher resolutions are split into two subclusters (13 and 15, Louv 1.6). We found upregulation of *Idgf* in cluster 9 (Louv 0.8, cluster 13, Louv 1.6); FISH revealed high expression in the terminal epithelium, and a lower expression level in the seminal vesicle. Cluster 10 (Louv 0.8) was enriched for *metallothionein A, methuselah-like 10* and *slimfast* (fig S9), all of which are enriched in terminal epithelium of *D. melanogaster* (Li, et al. 2022). We also found that *Ggt-1*, in addition to tail cyst cells, was expressed in the terminal epithelium, and enriched in both clusters 9 and 10 (fig 8F). Cluster 15 (Louv 1.6) was marked by high expression of *aqrs*, which was limited to the accessory glands (fig 8I).

#### The hub

The hub, a collection of 8-16 tightly packed somatic cells at the apical tip of the testis, maintains the germline and cyst stem cell populations, and contributes to stem cell differentiation by establishing signalling gradients (Hardy, et al. 1979; de Cuevas and Matunis 2011). FISH for the marker *esg* shows hub localisation (fig 3A) but was not upregulated in any individual cluster in the somatic scRNAseq data. Other hub markers *fas3* and *CadN* show upregulation in a subset of cluster 5 (Louv 0.8), although both are also markers of germline and cyst stem cells, and proliferating cells (fig S8).

#### The testis sheath: pigment and muscle cells

The testis wall is formed of three layers; pigment cells covering a layer of smooth multinucleated muscle, overlaying basement membrane. The muscle cells are difficult to stain by FISH, as they are extremely thin and cover the entire testis, resulting in low level ubiquitous signal, resembling background. We therefore inferred cluster identity of muscle marker *mhc*, observed in cluster 3 (Louv 0.8). Pigment cells are similarly difficult to image, although we were able to identify *pip*, which was upregulated in a subset of cluster 6 cells (Louv 0.8, cluster 18 Louv 3.2). FISH showed a low level of punctate signal surrounding the testis, consistent with pigment cells (fig S9).

#### Neurons

Cluster 11 (Louv 0.8) was enriched for multiple neuronal markers, including *VAchT, elav,* and *nSyb* (fig S9). Like the fly cell atlas data, we propose that a small number of neurons were captured during the dissection.

## Discussion

We have generated a whole-testis single cell RNA sequencing data set for a sperm heteromorphic species, *D. pseudoobscura*. As in the fly cell atlas dataset, germline and somatic cells clustered separately (Li, et al. 2022). The higher number of germline cells was, expected based on proportion of germline to somatic cells in testis (spermatocyte cysts comprise 32 germline cells compared to only two somatic cells) (Dobzhansky 1934; Kurokawa and Hihara 1976). We identified a diverse array of cell types and developmental stages, both in germline and somatic datasets, by a combination of previously identified marker genes and FISH. We were able to annotate germline developmental stages from early spermatogonia, through spermatocyte, meiosis, early spermatid to elongating spermatid stages. Crucially, we identified a split in the germline developmental trajectory and assigned eusperm and parasperm lineages based on differential gene expression. There was a second potential secondary trajectory split in the parasperm trajectory, consistent with the production of three sperm morphs in *D. pseudoobscura*.

### Eusperm and parasperm lineages are transcriptionally distinct from the spermatocyte stage, and are maintained throughout development

We identified a split in the germline developmental trajectory shortly after the spermatogonia to spermatocyte transition, which was maintained into the elongating spermatid stage. Cell fate specification therefore occurs no later than early spermatocytes but could occur earlier in development. Indeed, distinct populations of spermatogonia or stem cells may be present but are not detected in our dataset due to the smaller number and lower transcriptional activity of early cells.

### Provisioning and timing mechanisms may contribute to morph development

The early spermatocyte stage is associated with a marked increase in transcriptional activity – both in terms of transcriptome complexity, and in absolute transcript number (Fuller 1993; Raz, et al. 2023). Transcripts required for spermiogenesis are typically transcribed in spermatocytes and it is perhaps unsurprising that it is this stage where eusperm and parasperm transcriptional differences become obvious. Importantly, we found that euspermatocytes are more transcriptionally active than paraspermatocytes; euspermatocytes express more genes, and genes are more highly expressed. This finding indicates a provisioning mechanism in morph development - longer eusperm require more transcripts to make more proteins, from a greater number of genes.

During development, cysts are displaced away from the hub by subsequent divisions of the GSCs, moving basally in the radially symmetric testis. Cysts approximately the same distance from the hub should therefore be of a similar age. While examining expression patterns of germline-expressed genes, we noted differences in the positions of eu- and paraspermatocyte cysts within the testes. Euspermatocyte cysts were consistently positioned closer to the apical tip than stage-matched paraspermatocyte cysts. For example, *vasa* FISH showed a subset of relatively early spermatocyte cysts, with compact DNA, positioned amongst late spermatocyte and early spermatid cysts. Furthermore, *in situ* hybridisation of genes identified as differentially expressed between eu- and paraspermatocytes showed eusperm upregulated genes higher in somewhat larger, more apically positioned spermatocytes, which also had dimmer nuclear staining (less condensed DNA). Conversely, genes upregulated in parasperm were higher in somewhat smaller, more basally position spermatocyte cysts, which had brighter nuclear staining (more condensed DNA). We therefore suggest that paraspermatocytes mature more slowly than eusperm cysts.

Developmental timing, in concert with cyst positioning, may be important for morph specific development. Slower development of paraspermatocytes results in greater displacement of the paraspermatocyte cysts away from the hub, and towards the terminal epithelium. Post-meiotic paraspermatids will then have less distance, and therefore less time to elongate, before the heads reach the terminal epithelium, and individualisation is triggered.

### *TGIF* and *kmg* likely regulate transcriptional differences between eusperm and parasperm

We have documented extensive transcriptional differences between eusperm and parasperm, starting at the spermatocyte stage and continuing through at all subsequent stages of differentiation. To explore potential mechanisms for these differences, we characterised the expression of known testis transcription factors, finding eusperm upregulation of tMAC and TFIID components, *TGIF, wuc,* and *nht* and the testis-specific bromodomain-containing protein *tbrd-1*. Interestingly, we have previously identified a different component of tMAC to be implicated in heteromorphic sperm development in the greater waxmoth *Galleria mellonella* – the *aly* homologue *Lin9*, as well as the TFIID component *Taf4* (Moth, et al. 2024). While heteromorphic sperm has evolved independently in Lepidoptera and *Drosophila*, differentiation of heteromorphic sperm in these two lineages may share some common regulatory machinery.

TGIF is the *D. pseudoobscura* homologue of the *D. melanogaster* duplicate gene pair *achi* and *vis*, predicted to be DNA-binding subunits associated tMAC (Ayyar, et al. 2003; Wang and Mann 2003). In *D. melanogaster*, *achi/vis* and tMAC are required for transcription of essential genes in primary spermatocytes, binding to thousands of loci and acting as a pioneer transcription factor to open chromatin, enabling activation of spermatocyte specific promotors (Beall, et al. 2007; Matias, et al. 2025). All of these critical transcriptional regulators were expressed in both morphs indicating a conserved function in spermatogenic gene expression. However, *TGIF* transcript was upregulated, and maintained for longer, in euspermatocytes compared to paraspermatocytes, which indicates a potential role in the overall higher transcriptional activity we observed in this lineage. TGIF may more effectively recruit tMAC to the promotors of euspermatocyte specific genes as well as increasing transcription of target genes expressed in both morphs.

Conversely, we identified upregulation of the tMAC regulator *kmg* in paraspermatocytes. In *D. melanogaster* spermatocytes, Kmg maintains a germline transcriptional programme by recruiting chromatin modifiers Mi-2 and Dany to cryptic promotors of somatic and non-expressed genes, blocking tMAC and thus preventing inappropriate transcription (Kim, et al. 2017; Matias, et al. 2025). We did not detect significant upregulation of somatic transcripts in paraspermatocytes, indicating that cryptic promoters are effectively repressed in all early spermatocytes in *D. pseudoobscura*. *D. melanogaster* Kmg additionally works by counteracting tMAC at target promoters; at very highly expressed genes Kmg is displaced along the gene body from the promoters, while at more lowly expressed genes it remains associated with promoter regions (Kim, et al. 2017; Matias, et al. 2025). The higher level of Kmg protein in paraspermatocytes, and its greater chromatin localisation, is consistent with *D. pseudoobscura* Kmg acting at many tMAC target promoters tuning down transcription.

### Post-meiotic gene expression shows relatively little overlap with *D. melanogaster*, and shows euspermatid biased upregulation

While most genes whose products are required for spermiogenesis are transcribed in spermatocytes, a much smaller number (162 have been identified) are expressed after meiosis. Many of these mRNAs have a distinctive localisation pattern to the tail-end of elongating spermatids, hence their name ‘cup’ and ‘comet’ genes (Barreau, Benson, Gudmannsdottir, et al. 2008; Barreau, Benson and White-Cooper 2008; Raz, et al. 2023).

The functions of post-meiotically expressed genes are largely unknown. GO analysis of *D. melanogaster* suggests functions in transport, membranes, cytoskeleton, enzyme function and RNA regulation (Raz, et al. 2023). Functional studies of a few post-meiotic genes have been published; *scotti (soti)*, a post meiotic gene in *D. melanogaster,* regulates individualisation by inhibiting caspase activation in individualising spermatids (Kaplan, et al. 2010). The post-meiotically transcribed RNA-binding protein Imp binds to multiple post-meiotic transcripts, including *world-cup, soti, whip, schuy* and *c-cup*, and may have a role in regulating subcellular localisation of these transcripts (Jackson 2024). Given how little is understood about the roles of post-meiotically transcribed genes in *D. melanogaster*, it is difficult to hypothesise potential roles in heteromorphic sperm development, but they may contribute to the timing of elongation and individualisation, transport and translation of sperm tail components, or are components of mature sperm.

It was notable that relatively few genes previously identified as post-meiotically transcribed in *D. melanogaster* were also identified in *D. pseudoobscura*; of 203 *D. pseudoobscura* post-meiotic genes only were also post-meiotic in *D. melanogaster* genes. One particularly surprising example was *comr*, a component of tMAC which is not post-meiotically transcribed in *D. melanogaster* but is transcribed in *D. pseudoobscura* post-meiotic spermatids, producing an mRNA localised to spermatid distal tail-ends. Future work could capitalise on this apparent lack of conservation to explore the mechanisms by which genes gain or lose post-meiotic transcription.

We also found a bias towards post-meiotic genes being upregulated in euspermatids, compared to paraspermatids. Post-meiotically transcribed genes were either detected equally in both morphs, or were upregulated in euspermatids. We did not identify any genes upregulated in paraspermatids compared to euspermatids. This would be consistent with a role for such transcripts in eusperm elongation and timing of individualisation.

### Gene duplication is a mechanism for morph-specific gene specialisation

We identified several examples of duplicated genes pairs gaining differential gene expression, which may indicate a morph-specific or morph-enriched gene function. One such example was that of the protamines, which replace nucleosomes in spermatid chromatin. While *protamine A* was expressed equally in both morphs, *Protamine-like SSC* and *protamine* were differentially expressed in eu- and paraspermatocytes. Parasperm nuclei are shorter and wider than eusperm nuclei; potentially alterations to DNA packaging between the morphs could underly this gross morphological difference.

A more extreme example was that of prominin-like protein family, which has 13 paralogues in *D. pseudoobscura*, compared to two in *D. melanogaster*. Of these 13, two were not detected in testes, five were expressed in both morphs but not differentially expressed, and six were differentially expressed. More promL proteins were upregulated in the eusperm trajectory, which may reflect requirement for higher levels of promL proteins in longer sperm, or that eusperm requires additional functions which are carried out by these paralogues. In two instances we found that the most recently duplication event had generated a pair of genes with upregulation in different morphs. It is likely that the duplication of these genes facilitated in morph-specific subfunctionalisation.

Testis-biased expression of new genes is a well-described phenomenon (Levine, et al. 2006; White-Cooper and Bausek 2010; Assis and Bachtrog 2013; Kondo, et al. 2017; Rivard, et al. 2021). Sperm heteromorphy may present an additional opportunity for neofunctionalization within the testis. Young genes which gain testis-expression via the permissive transcriptional environment of chromatin in the male germline may acquire morph-specific regulatory elements. Gene duplicates can then be expressed in one morph, without their functions affecting the other, and are exposed to selective pressures resulting in morph-specific or morph-enriched neofunctionalization.

### Parasperm morphs not clear from transcriptional data

The ancestral state of sperm morphology in *Drosophila* is monomorphic; the derived state in the *obscura* species group is dimorphic, with the *pseudoobscura* sub-group having secondarily evolved trimorphic sperm (Messer and White-Cooper 2025).

We have previously confirmed the presence of two parasperm morphs in *D. pseudoobscura* using the SLoB3 line also used in this study (Messer and White-Cooper 2025). Parasperm morphs have distinct lengths: Parasperm 1 are approximately 45μm with a 10μm nucleus, whereas parasperm 2 are approximately 90μm, with a 15μm nucleus. It is perhaps unexpected then, that we did not observe a dramatic split in the parasperm developmental trajectory, which could be assigned to parasperm 1 and 2 morphs. At higher clustering resolution, we were able to identify a split in the mid-paraspermatocyte stage, but this was not maintained through the later trajectory. This suggests that parasperm 1 and 2 are transcriptionally very similar, although a larger dataset may be able to better distinguish the parasperm 1 and 2 developmental trajectories. Nevertheless, the evolutionarily most recent innovation, the parasperm split, occurs after the than eusperm-parasperm identity specification.

### No evidence for morph-specific cyst cell populations

Somatic cyst cells have essential functions in regulating the environment of the developing germline cells, including transporting metabolites and nutrients, and signalling to regulate key germline developmental transitions (Matunis, et al. 1997; Hudry, et al. 2019; Messer, et al. 2026; Sainz de la Maza, et al. 2026). We were able to identify head and tail cyst cell populations associated with elongating spermatids, as has previously been observed in *D. melanogaster* (Raz, et al. 2023). Head and tail cyst cells are morphologically distinct; head cyst cells are compact, covering the heads of the spermatid bundle, whereas tail cyst cells are extremely elongated to cover the spermatid tails (Tokuyasu, et al. 1972). Head cyst cells associate with the terminal epithelium to initiate spermatid coiling and release from the cyst (Tokuyasu, et al. 1972).

While we found clear evidence of eusperm and parasperm developmental trajectories in the germline data, we did not find any evidence of multiple somatic cyst cell populations which specifically associate with eusperm or parasperm cysts, either in the scRNAseq data or any of our FISH experiments. If there are distinct cyst cell populations the differences are likely extremely subtle or enacted at the translational level. We therefore suggest that the regulation of sperm length is likely germline-intrinsic, as recently shown via germ cell transplantation in *D. simulans* (Nishimura, et al. 2026).

### Provisioning and timing factors are implicated in determining cell length

We hypothesised four mechanisms by which sperm length may be determined in sperm heteromorphic *Drosophila*: provisioning, timing, blocking and measuring. We found strong evidence that provisioning and timing have roles in sperm length, as indicated by differences we identified between developing eusperm and parasperm. The parasperm lineage progresses through differentiation more slowly than eusperm, suggesting that timing of key developmental transitions may contribute to regulating sperm length, perhaps in concert with the terminal epithelium providing a stop signal. Developing eusperm express more genes, and have higher transcription than parasperm, i.e. they have more provisions.

## Supplementary Files

1. Dpse_germline.loom - Loom file of germline scRNAseq dataset
2. Dpse_somatic.loom - Loom file of somatic scRNAseq dataset
3. Dpse_testis.loom – Loom file of whole testis scRNAseq dataset
4. piggyBac-Kmg-GFP.gb – genbank sequence file for piggyBac Kmg::GFP fusion construct
5. Supplementary methods
6. Supplementary figures 1-6, 8, 9
7. supp_fig_7_Dpse_somatic3D.html - Supplementary figure 7
8. Supplementary tables 1-6

## Acknowledgements

We are especially grateful to Tom Price from the University of Liverpool for giving us his *D. pseudoobscura* stocks, and to the Genome, Biocomputing and Bioimaging (RRID: SCR_022556) hubs in the School of Biosciences at Cardiff University for access to facilities and their technical support. We thank colleagues in the Drosophila community, in particular FlyBase and the National Drosophila Species Stock Center NDSSC). We are grateful to many colleagues including in the Cardiff fly community for their critical discussions and feedback on this manuscript. We thank former lab members Jasmine Caldwell, Daniel Michael and Emma Moth for their contribution to pilot data for this work. This work was funded by Leverhulme Trust project grant RPG-2023-195 and the BBSRC South West Biosciences Doctoral Training Partnership BB/M009122/1.

## Data Availability

Raw sequence data is available via NCBI Sequence Read Archive, BioProject PRJNA1497681.

## Notes

### Competing Interest Statement

The authors have declared no competing interest.

## References

1. Alpern JHM, Asselin MM, Moehring AJ. 2019. Identification of a novel sperm class and its role in fertilization in *Drosophila*. J Evol Biol 32:259–266.

2. Altschul SF, Madden TL, Schaffer AA, Zhang J, Zhang Z, Miller W, Lipman DJ. 1997. Gapped BLAST and PSI-BLAST: a new generation of protein database search programs. Nucleic Acids Res 25:3389–3402.

3. FastQC – A Quality Control tool for High Throughput Sequence Data. [Internet]. Babraham Bioinformatics; 2010 [cited 2017 November 20]. Available from: http://www.bioinformatics.babraham.ac.uk/projects/fastqc/

4. Assis R, Bachtrog D. 2013. Neofunctionalization of young duplicate genes in *Drosophila*. Proc Natl Acad Sci U S A 110:17409–17414.

5. Ayyar S, Jiang J, Collu A, White-Cooper H, White RA. 2003. *Drosophila* TGIF is essential for developmentally regulated transcription in spermatogenesis. Development 130:2841–2852.

6. Barreau C, Benson E, Gudmannsdottir E, Newton F, White-Cooper H. 2008. Post-meiotic transcription in *Drosophila* testes. Development 135:1897–1902.

7. Barreau C, Benson E, White-Cooper H. 2008. Comet and cup genes in *Drosophila* spermatogenesis: the first demonstration of post-meiotic transcription. Biochem Soc Trans 36:540–542.

8. Beall EL, Lewis PW, Bell M, Rocha M, Jones DL, Botchan MR. 2007. Discovery of tMAC: a *Drosophila* testis-specific meiotic arrest complex paralogous to Myb-Muv B. Genes Dev 21:904– 919.

9. Beatty RA, Sidhu NS. 1970. Polymegaly of spermatozoan length and its genetic control in *Drosophila* species. Proceedings of the Royal Society of Edinburgh Section B: Biological Sciences 71B:14–29.

10. Bhatt K. 2025. Novel Origin of Sperm Heteromorphism in the Drosophilidae. [M.S.]. [United States -- New York]: Syracuse University.

11. Choi HMT, Schwarzkopf M, Fornace ME, Acharya A, Artavanis G, Stegmaier J, Cunha A, Pierce NA. 2018. Third-generation in situ hybridization chain reaction: multiplexed, quantitative, sensitive, versatile, robust. Development 145.

12. de Cuevas M, Matunis EL. 2011. The stem cell niche: lessons from the *Drosophila* testis. Development 138:2861–2869.

13. Dobin A, Davis CA, Schlesinger F, Drenkow J, Zaleski C, Jha S, Batut P, Chaisson M, Gingeras TR. 2013. STAR: ultrafast universal RNA-seq aligner. Bioinformatics 29:15–21.

14. Dobzhansky T. 1934. Studies on hybrid sterility. Z Zellforsch Mikrosk Anat 21:169–223.

15. Fabian L, Brill JA. 2012. *Drosophila* spermiogenesis: Big things come from little packages. Spermatogenesis 2:197–212.

16. Fabrizio JJ, Boyle M, DiNardo S. 2003. A somatic role for eyes absent (eya) and sine oculis (so) in Drosophila spermatocyte development. Dev Biol 258:117–128.

17. Fargeas CA, Joester A, Missol-Kolka E, Hellwig A, Huttner WB, Corbeil D. 2004. Identification of novel Prominin-1/CD133 splice variants with alternative C-termini and their expression in epididymis and testis. J Cell Sci 117:4301–4311.

18. Fuller M. 1993. Spermatogenesis. In: Bate M, Martinez Arias A, editors. Development of Drosophila: Cold Spring Harbor Laboratory Press. p. 71–147.

19. Han YG, Kwok BH, Kernan MJ. 2003. Intraflagellar transport is required in Drosophila to differentiate sensory cilia but not sperm. Curr Biol 13:1679–1686.

20. Handler AM, McCombs SD, Fraser MJ, Saul SH. 1998. The lepidopteran transposon vector, *piggyBac*, mediates germ-line transformation in the Mediterranean fruit fly. Proc Natl Acad Sci U S A 95:7520–7525.

21. Hao Y, Stuart T, Kowalski MH, Choudhary S, Hoffman P, Hartman A, Srivastava A, Molla G, Madad S, Fernandez-Granda C, et al. 2024. Dictionary learning for integrative, multimodal and scalable single-cell analysis. Nat Biotechnol 42:293–304.

22. Hardy RW, Tokuyasu KT, Lindsley DL, Garavito M. 1979. The germinal proliferation center in the testis of *Drosophila melanogaster*. J Ultrastruct Res 69:180–190.

23. Hiller M, Chen X, Pringle MJ, Suchorolski M, Sancak Y, Viswanathan S, Bolival B, Lin TY, Marino S, Fuller MT. 2004. Testis-specific TAF homologs collaborate to control a tissue-specific transcription program. Development 131:5297–5308.

24. Hiller MA, Lin TY, Wood C, Fuller MT. 2001. Developmental regulation of transcription by a tissue-specific TAF homolog. Genes Dev 15:1021–1030.

25. Holman L, Snook RR. 2008. A sterile sperm caste protects brother fertile sperm from female-mediated death in *Drosophila pseudoobscura*. Curr Biol 18:292–296.

26. Holtzman S, Miller D, Eisman R, Kuwayama H, Niimi T, Kaufman T. 2010. Transgenic tools for members of the genus *Drosophila* with sequenced genomes. Fly (Austin) 4:349–362.

27. Hudry B, de Goeij E, Mineo A, Gaspar P, Hadjieconomou D, Studd C, Mokochinski JB, Kramer HB, Placais PY, Preat T, et al. 2019. Sex Differences in Intestinal Carbohydrate Metabolism Promote Food Intake and Sperm Maturation. Cell 178:901–918 e916.

28. Hurbain I, Mace AS, Romao M, Prince E, Sengmanivong L, Ruel L, Basto R, Therond PP, Raposo G, D’Angelo G. 2022. Microvilli-derived extracellular vesicles carry Hedgehog morphogenic signals for Drosophila wing imaginal disc development. Curr Biol 32:361–373 e366.

29. Jackson D. 2024. RNA binding proteins and mRNA localisation in Drosophila sperm development. [Cardiff University.

30. Jiang J, White-Cooper H. 2003. Transcriptional activation in *Drosophila* spermatogenesis involves the mutually dependent function of *aly* and a novel meiotic arrest gene *cookie monster*. Development 130:563–573.

31. Kaplan Y, Gibbs-Bar L, Kalifa Y, Feinstein-Rotkopf Y, Arama E. 2010. Gradients of a ubiquitin E3 ligase inhibitor and a caspase inhibitor determine differentiation or death in spermatids. Dev Cell 19:160–173.

32. Katzenberger RJ, Rach EA, Anderson AK, Ohler U, Wassarman DA. 2012. The Drosophila Translational Control Element (TCE) is required for high-level transcription of many genes that are specifically expressed in testes. PLoS One 7:e45009.

33. Kim J, Lu C, Srinivasan S, Awe S, Brehm A, Fuller MT. 2017. Blocking promiscuous activation at cryptic promoters directs cell type-specific gene expression. Science 356:717–721.

34. Kondo S, Vedanayagam J, Mohammed J, Eizadshenass S, Kan L, Pang N, Aradhya R, Siepel A, Steinhauer J, Lai EC. 2017. New genes often acquire male-specific functions but rarely become essential in *Drosophila*. Genes Dev 31:1841–1846.

35. Kurokawa H, Hihara F. 1976. Number of first spermatocytes in relation to phylogeny of Drosophila (Diptera : Drosophilidae). International Journal of Insect Morphology and Embryology 5:51–63.

36. Laktionov PP, Maksimov DA, Romanov SE, Antoshina PA, Posukh OV, White-Cooper H, Koryakov DE, Belyakin SN. 2018. Genome-wide analysis of gene regulation mechanisms during *Drosophila* spermatogenesis. Epigenetics Chromatin 11:14.

37. Laktionov PP, White-Cooper H, Maksimov DA, Beliakin SN. 2014. [Transcription factor comr acts as a direct activator in the genetic program controlling spermatogenesis in D. melanogaster]. Mol Biol (Mosk) 48:153–165.

38. Lee E, Sivan-Loukianova E, Eberl DF, Kernan MJ. 2008. An IFT-A protein is required to delimit functionally distinct zones in mechanosensory cilia. Curr Biol 18:1899–1906.

39. Levine MT, Jones CD, Kern AD, Lindfors HA, Begun DJ. 2006. Novel genes derived from noncoding DNA in *Drosophila melanogaster* are frequently X-linked and exhibit testis-biased expression. Proc Natl Acad Sci U S A 103:9935–9939.

40. Li H, Handsaker B, Wysoker A, Fennell T, Ruan J, Homer N, Marth G, Abecasis G, Durbin R, Subgroup GPDP. 2009. The Sequence alignment/map (SAM) format and SAMtools. Bioinformatics 25:2078–2079.

41. Li H, Janssens J, De Waegeneer M, Kolluru SS, Davie K, Gardeux V, Saelens W, David FPA, Brbic M, Spanier K, et al. 2022. Fly Cell Atlas: A single-nucleus transcriptomic atlas of the adult fruit fly. Science 375:eabk2432.

42. Lindsley DL, Tokuyasu KT. 1980. Spermatogenesis. In: Ashburner M, F. Wtr, editors. The Genetics and Biology of Drosophila. London: Academic Press. p. 225–294.

43. Lu D, Sin HS, Lu C, Fuller MT. 2020. Developmental regulation of cell type-specific transcription by novel promoter-proximal sequence elements. Genes Dev 34:663–677.

44. Machlab D, Burger L, Soneson C, Rijli FM, Schubeler D, Stadler MB. 2022. monaLisa: an R/Bioconductor package for identifying regulatory motifs. Bioinformatics 38:2624–2625.

45. Madeira F, Madhusoodanan N, Lee J, Eusebi A, Niewielska A, Tivey ARN, Lopez R, Butcher S. 2024. The EMBL-EBI Job Dispatcher sequence analysis tools framework in 2024. Nucleic Acids Res 52:W521–W525.

46. Mahato S, Nie J, Plachetzki DC, Zelhof AC. 2018. A mosaic of independent innovations involving eyes shut are critical for the evolutionary transition from fused to open rhabdoms. Dev Biol 443:188–202.

47. Martinez-Campos M, Basto R, Baker J, Kernan M, Raff JW. 2004. The Drosophila pericentrin-like protein is essential for cilia/flagella function, but appears to be dispensable for mitosis. J Cell Biol 165:673–683.

48. Matias NR, Gallicchio L, Lu D, Kim JJ, Perez J, Detweiler AM, Lu C, Bolival B, Fuller MT. 2025. A cell type-specific surveillance complex represses cryptic promoters during differentiation in an adult stem cell lineage. Genes Dev 39:1318–1337.

49. Matsukuma H, Kobayashi Y, Oka S, Higashijima F, Kimura K, Yoshihara E, Sasai N, Shiraishi K. 2023. Prominin-1 deletion results in spermatogenic impairment, sperm morphological defects, and infertility in mice. Reprod Med Biol 22:e12514.

50. Matunis E, Tran J, Gonczy P, Caldwell K, DiNardo S. 1997. punt and schnurri regulate a somatically derived signal that restricts proliferation of committed progenitors in the germline. Development 124:4383–4391.

51. Messer F, White-Cooper H, Amoyel M. 2026. Spermatogenesis. In: Yamanaka N, Atkinson PW, editors. Comprehensive Molecular Insect Science. Oxford: Elsevier. p. 356–390.

52. Messer F, White-Cooper H. 2025. Clustering sperm: A statistical approach to identify sperm morph numbers in the Drosophila obscura species group. Physiological Entomology 51:48–62.

53. Messer FR. 2022. Developmental and molecular analysis of sperm in Drosophila pseudoobscura. [Cardiff University.

54. Metcalf CE, Wassarman DA. 2007. Nucleolar colocalization of TAF1 and testis-specific TAFs during *Drosophila* spermatogenesis. Dev Dyn 236:2836–2843.

55. Moore AJ, Bacigalupe LD, Snook RR. 2013. Integrated and independent evolution of heteromorphic sperm types. Proc Biol Sci 280:20131647.

56. Moth E, Messer F, Chaudhary S, White-Cooper H. 2024. Differential gene expression underpinning the production of distinct sperm morphs in the wax moth Galleria mellonella. Open Biol 14:240002.

57. Navarro Paya D. 2017. Gene drive in Drosophila melanogaster and Aedes aegypti. [Cardiff University.

58. Nishimura K, Asaoka M, Takano-Shimizu-Kouno T. 2026. Germline and male somatic determinants of sperm length divergence in Drosophila: implications for postcopulatory sexual selection. Genetics 233:iyag069.

59. Olivieri G, Olivieri A. 1965. Autoradiographic study of nucleic acid synthesis during spermatogenesis in *Drosophila melanogaster*. Mutat Res 2:366–380.

60. Pasini ME, Redi CA, Caviglia O, Perotti ME. 1996. Ultrastructural and cytochemical analysis of sperm dimorphism in *Drosophila subobscura*. Tissue Cell 28:165–175.

61. Policansky D, Ellison J. 1970. “Sex ratio” in Drosophila pseudoobscura: spermiogenic failure. Science 169:888–889.

62. Presgraves DC, Baker RH, Wilkinson GS. 1999. Coevolution of sperm and female reproductive tract morphology in stalk–eyed flies. Proceedings of the Royal Society of London. Series B: Biological Sciences 266:1041–1047.

63. Price MN, Dehal PS, Arkin AP. 2010. FastTree 2--approximately maximum-likelihood trees for large alignments. PLoS One 5:e9490.

64. R: A Language and Environment for Statistical Computing [Internet]. Vienna, Austria: R Foundation for Statistical Computing; 2025. Available from: https://www.R-project.org/

65. Rauluseviciute I, Riudavets-Puig R, Blanc-Mathieu R, Castro-Mondragon JA, Ferenc K, Kumar V, Lemma RB, Lucas J, Cheneby J, Baranasic D, et al. 2024. JASPAR 2024: 20th anniversary of the open-access database of transcription factor binding profiles. Nucleic Acids Res 52:D174–D182.

66. Raz AA, Vida GS, Stern SR, Mahadevaraju S, Fingerhut JM, Viveiros JM, Pal S, Grey JR, Grace MR, Berry CW, et al. 2023. Emergent dynamics of adult stem cell lineages from single nucleus and single cell RNA-Seq of Drosophila testes. eLife 12:e82201.

67. Richens JH, Barros TP, Lucas EP, Peel N, Pinto DM, Wainman A, Raff JW. 2015. The Drosophila Pericentrin-like-protein (PLP) cooperates with Cnn to maintain the integrity of the outer PCM. Biol Open 4:1052–1061.

68. Rivard EL, Ludwig AG, Patel PH, Grandchamp A, Arnold SE, Berger A, Scott EM, Kelly BJ, Mascha GC, Bornberg-Bauer E, et al. 2021. A putative de novo evolved gene required for spermatid chromatin condensation in Drosophila melanogaster. PLoS Genet 17:e1009787.

69. Ryu TH, Subramanian M, Yeom E, Yu K. 2022. The prominin-like Gene Expressed in a Subset of Dopaminergic Neurons Regulates Locomotion in Drosophila. Mol Cells 45:640–648.

70. Ryu TH, Yeom E, Subramanian M, Lee KS, Yu K. 2019. Prominin-like Regulates Longevity and Glucose Metabolism via Insulin Signaling in Drosophila. J Gerontol A Biol Sci Med Sci 74:1557– 1563.

71. Sainz de la Maza D, Jefferson H, Brucker CI, Paoli S, Amoyel M. 2026. Somatic cells compartmentalise their carbohydrate metabolism to sustain germ cell survival. EMBO J.

72. Schafer M, Nayernia K, Engel W, Schafer U. 1995. Translational control in spermatogenesis. Dev Biol 172:344–352.

73. Schaffer AA, Aravind L, Madden TL, Shavirin S, Spouge JL, Wolf YI, Koonin EV, Altschul SF. 2001. Improving the accuracy of PSI-BLAST protein database searches with composition-based statistics and other refinements. Nucleic Acids Res 29:2994–3005.

74. Shao Z, Hu J, Jandura A, Wilk R, Jachimowicz M, Ma L, Hu C, Sundquist A, Das I, Samuel-Larbi P, et al. 2024. Spatially revealed roles for lncRNAs in Drosophila spermatogenesis, Y chromosome function and evolution. Nat Commun 15:3806.

75. Snook RR, Karr TL. 1998. Only long sperm are fertilization-competent in six sperm-heteromorphic *Drosophila* species. Curr Biol 8:291–294.

76. Snook RR, Markow TA, Karr TL. 1994. Functional nonequivalence of sperm in *Drosophila pseudoobscura*. Proc Natl Acad Sci U S A 91:11222–11226.

77. Swallow JG, Wilkinson GS. 2002. The long and short of sperm polymorphisms in insects. Biol Rev Camb Philos Soc 77:153–182.

78. Theofel I, Bartkuhn M, Hundertmark T, Boettger T, Gartner SM, Leser K, Awe S, Schipper M, Renkawitz-Pohl R, Rathke C. 2014. tBRD-1 selectively controls gene activity in the *Drosophila* testis and interacts with two new members of the bromodomain and extra-terminal (BET) family. PLoS One 9:e108267.

79. Tokuyasu KT, Peacock WJ, Hardy RW. 1972. Dynamics of spermiogenesis in Drosophila melanogaster. II. Coiling process. Z Zellforsch Mikrosk Anat 127:492–525.

80. Wang X, Zheng H, Jia Z, Lei Z, Li M, Zhuang Q, Zhou H, Qiu Y, Fu Y, Yang X, et al. 2019. Drosophila Prominin-like, a homolog of CD133, interacts with ND20 to maintain mitochondrial function. Cell Biosci 9:101.

81. Wang Z, Mann RS. 2003. Requirement for two nearly identical TGIF-related homeobox genes in *Drosophila* spermatogenesis. Development 130:2853–2865.

82. White-Cooper H, Bausek N. 2010. Evolution and spermatogenesis. Philos Trans R Soc Lond B Biol Sci 365:1465–1480.

83. White-Cooper H, Davidson I. 2011. Unique aspects of transcription regulation in male germ cells. Cold Spring Harb Perspect Biol 3.

84. White-Cooper H, Leroy D, MacQueen A, Fuller MT. 2000. Transcription of meiotic cell cycle and terminal differentiation genes depends on a conserved chromatin associated protein, whose nuclear localisation is regulated. Development 127:5463–5473.

85. Wingert L, DiNardo S. 2015. Traffic jam functions in a branched pathway from Notch activation to niche cell fate. Development 142:2268–2277.

86. Zheng H, Zhang Y, Chen Y, Guo P, Wang X, Yuan X, Ge W, Yang R, Yan Q, Yang X, et al. 2019. Prominin-like, a homolog of mammalian CD133, suppresses di lp6 and TOR signaling to maintain body size and weight in Drosophila. FASEB J 33:2646–2658.

87. Zur Lage P, Newton FG, Jarman AP. 2019. Survey of the Ciliary Motility Machinery of *Drosophila* Sperm and Ciliated Mechanosensory Neurons Reveals Unexpected Cell-Type Specific Variations: A Model for Motile Ciliopathies. Front Genet 10:24.

