## Supplementary material for "Single cell RNA sequencing of *D. pseudoobscura* testes reveals transcriptional signatures of heteromorphic spermatogenesis": Supplementary figures.pdf

**Figure S1**

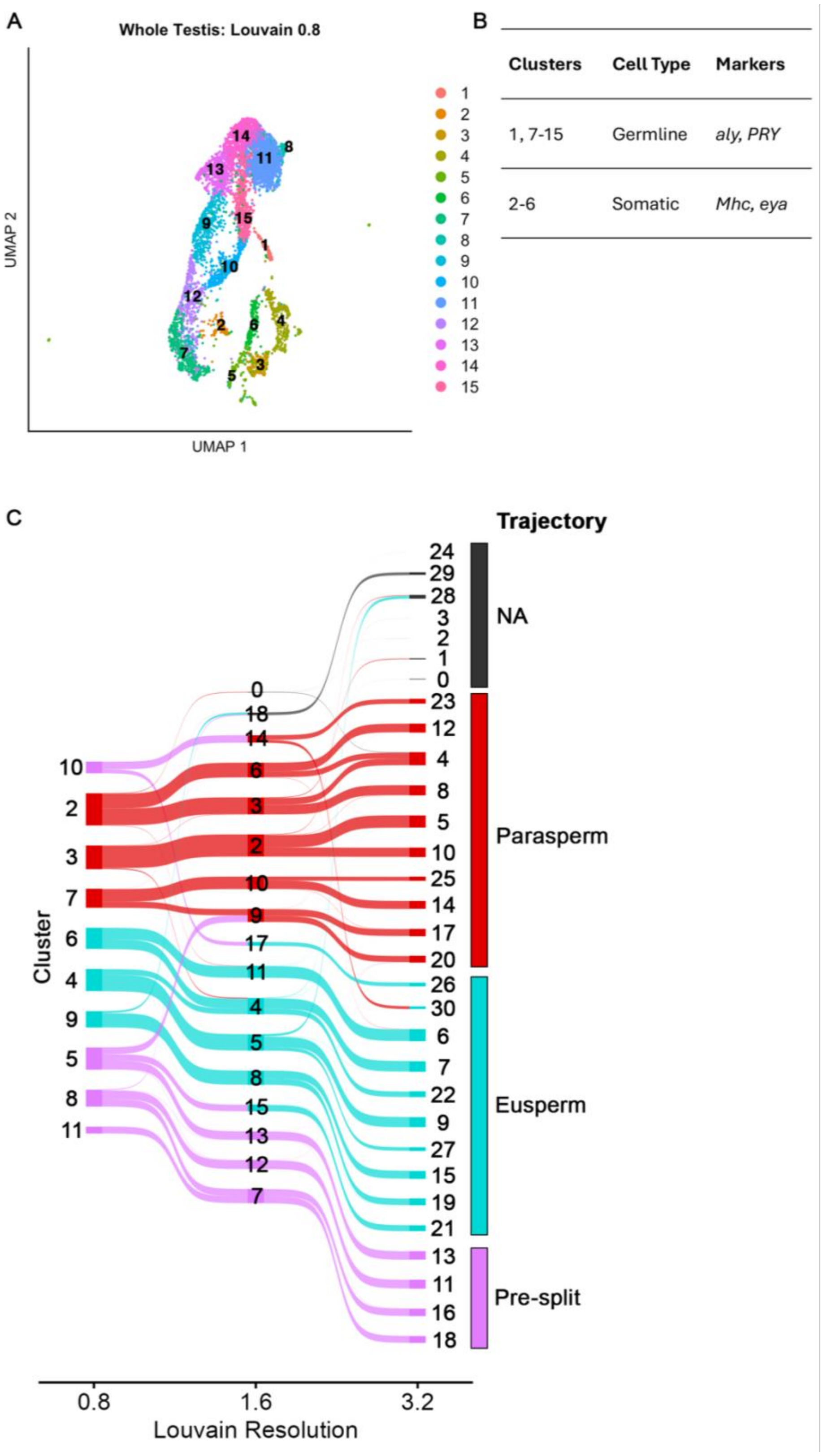


| Clusters | Cell Type | Markers |
|---|---|---|
| 1, 7-15 | Germline | *aly, PRY* |
| 2-6 | Somatic | *Mhc, eya* |

**Supplementary figure 1: Summary of clustering in whole testis and germline datasets.** (A) UMAP of whole testis scRNAseq, with Louvain clustering at resolution 0.8.

(B) Table of whole testis clusters assigned to germline and somatic datasets. (C) Sankey diagram of germline clustering at resolutions 0.8, 1.6 and 3.2, showing division of low-resolution clusters into subclusters at higher resolutions. Node width indicates cluster size, flow width indicates the proportion of cells in a given cluster contributing to higher resolution clusters. Trajectory identity is indicated by node and flow colour. NA indicates no assignment of that cluster to a trajectory. Low levels of cross-cluster branching indicate high cluster stability.

**Figure S2**

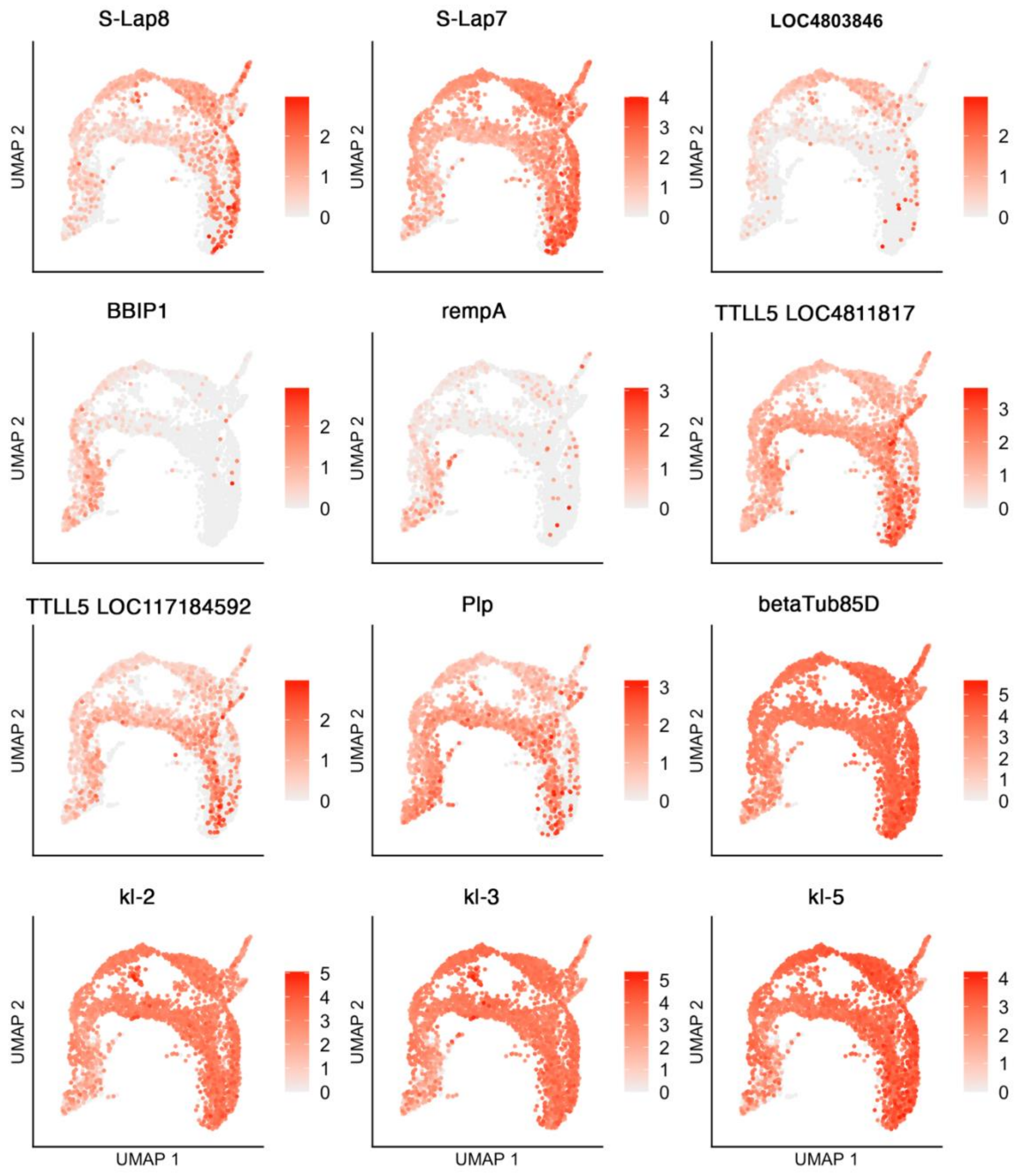


**Supplementary figure 2: Germline UMAPs of differentially expressed flagellar assembly and sperm tail components.** S-Lap8 and S-Lap7 are components of the paracrystalline material in the mitochondrial derivates of mature sperm. *LOC4803846* is orthologous to *CG30268*, predicted to encode a component of the axoneme radial spoke. BBIP1 and rempA function in cilium assembly and transport. TTLL5 is a tubulin-tyrosine ligase, although its function in sperm is unknown. Plp is a component of the pericentriolar material. βTub85D is then sperm axoneme beta tubulin. Kl-2, Kl-3 and Kl-5 are sperm tail kinesins.

**Figure S3**

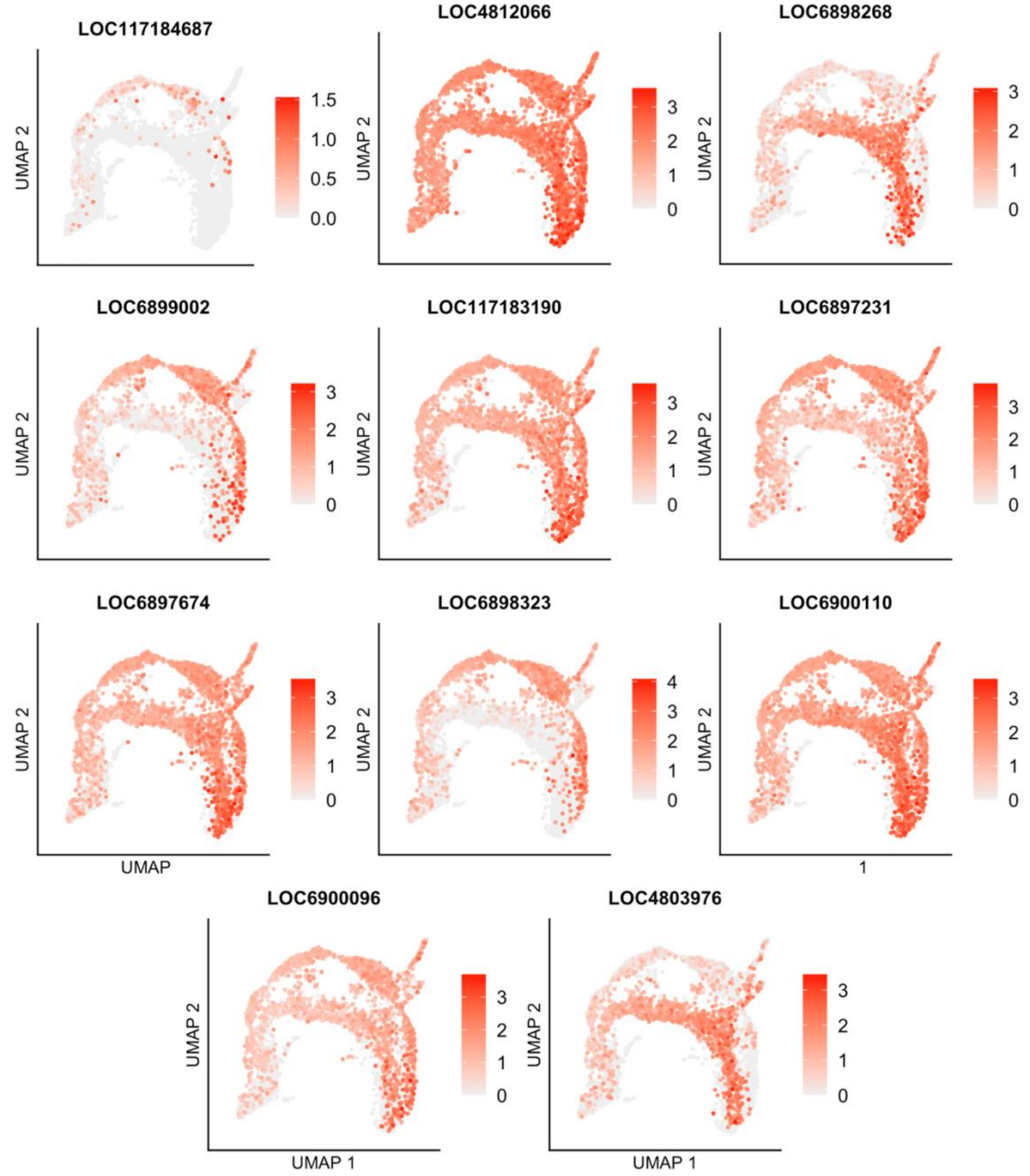


**Supplementary figure 3: Expression UMAPs of *D. pseudoobscura* prominin-like genes in germline.**

**Figure S4**

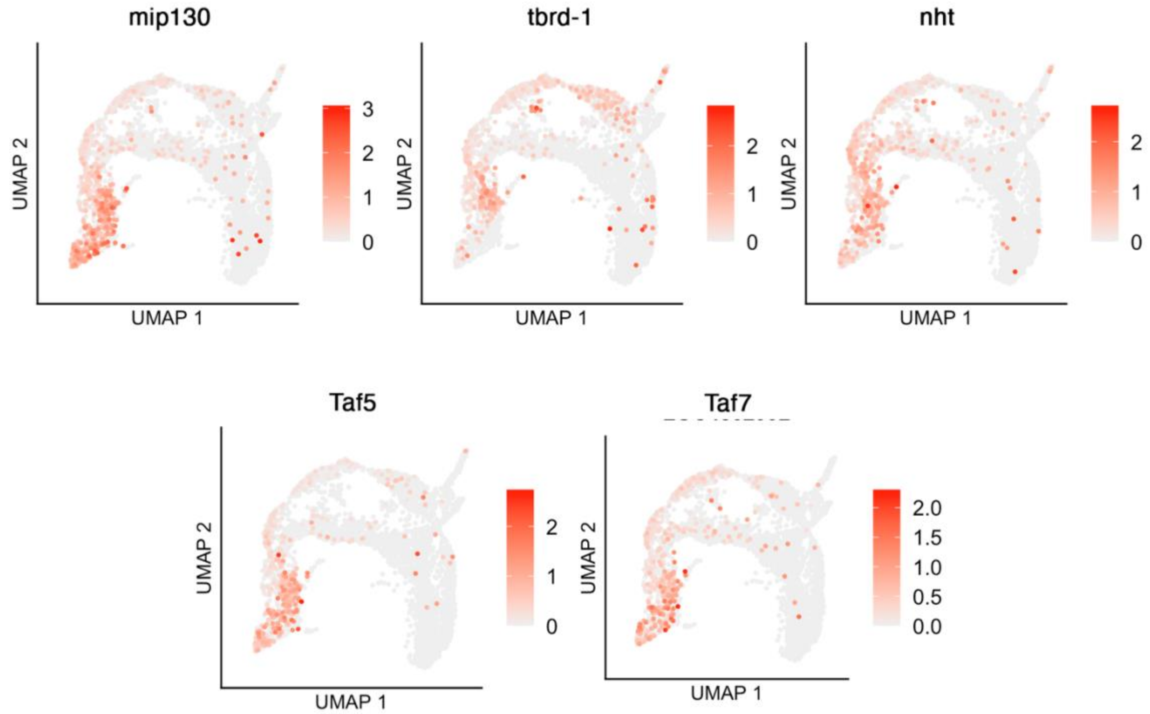


**Supplementary figure 4: Expression UMAPs of transcription factor genes.** Mip130 is the somatic paralogue of *aly* in *D. melanogaster*. Tbrd-1 is a testis-specific bromodomain containing protein. Nht is a component of the predicted testis specific paralogue of the TFIID transcription complex, tTFIID. Taf5 and Taf7 are components of TFIID.

**Figure S5**

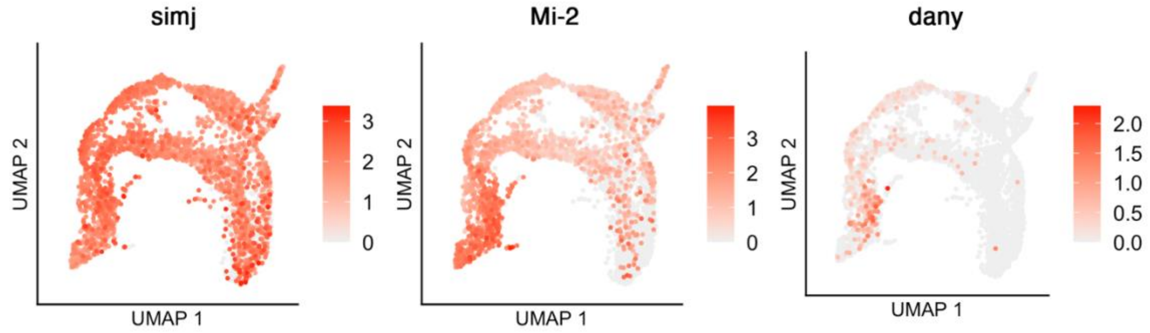


**Supplementary figure 5: Expression UMAPs of Kmg-interacting proteins.**

**Figure S6**

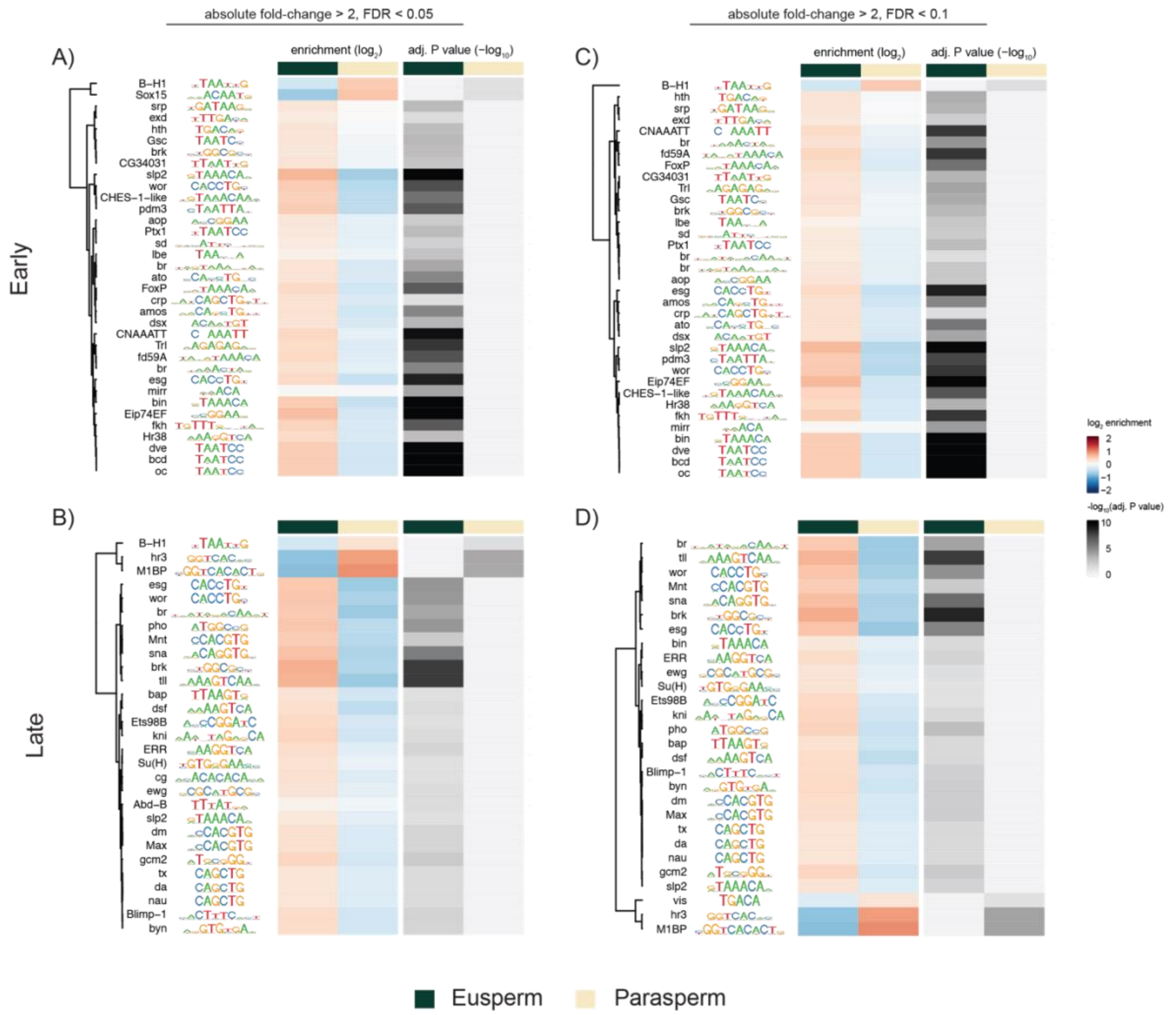


**Supplementary figure 6: Motif enrichment analysis implicates distinct sets of TFs in eusperm and parasperm specific expression.** (A-B) Heatmap displaying log2 enrichment and adjusted p-values for motifs present in genes identified as differentially expressed (absolute FC > 2, FDR < 0.05) at (A) early and (B) late timepoints. Heatmap displaying log2 enrichment and adjusted p-values for motifs present in genes identified as differentially expressed (absolute FC > 2, FDR < 0.1) at early (C) and late (D) timepoints. (A, C) Comparison between euspermatocyte cluster 21 and paraspermatocyte cluster 20 (Louvain 3.2). (B, D) Comparison between mid-euspermatocyte cluster 9 and mid-paraspermatocyte cluster 7 (Louvain 0.8).

**Figure S7**

supp_fig_7_Dpse_somatic3D.html

**Supplementary figure 7: 3D cluster UMAP of somatic cell data, clustered at Louvain resolution 0.8.**

**Figure S8**

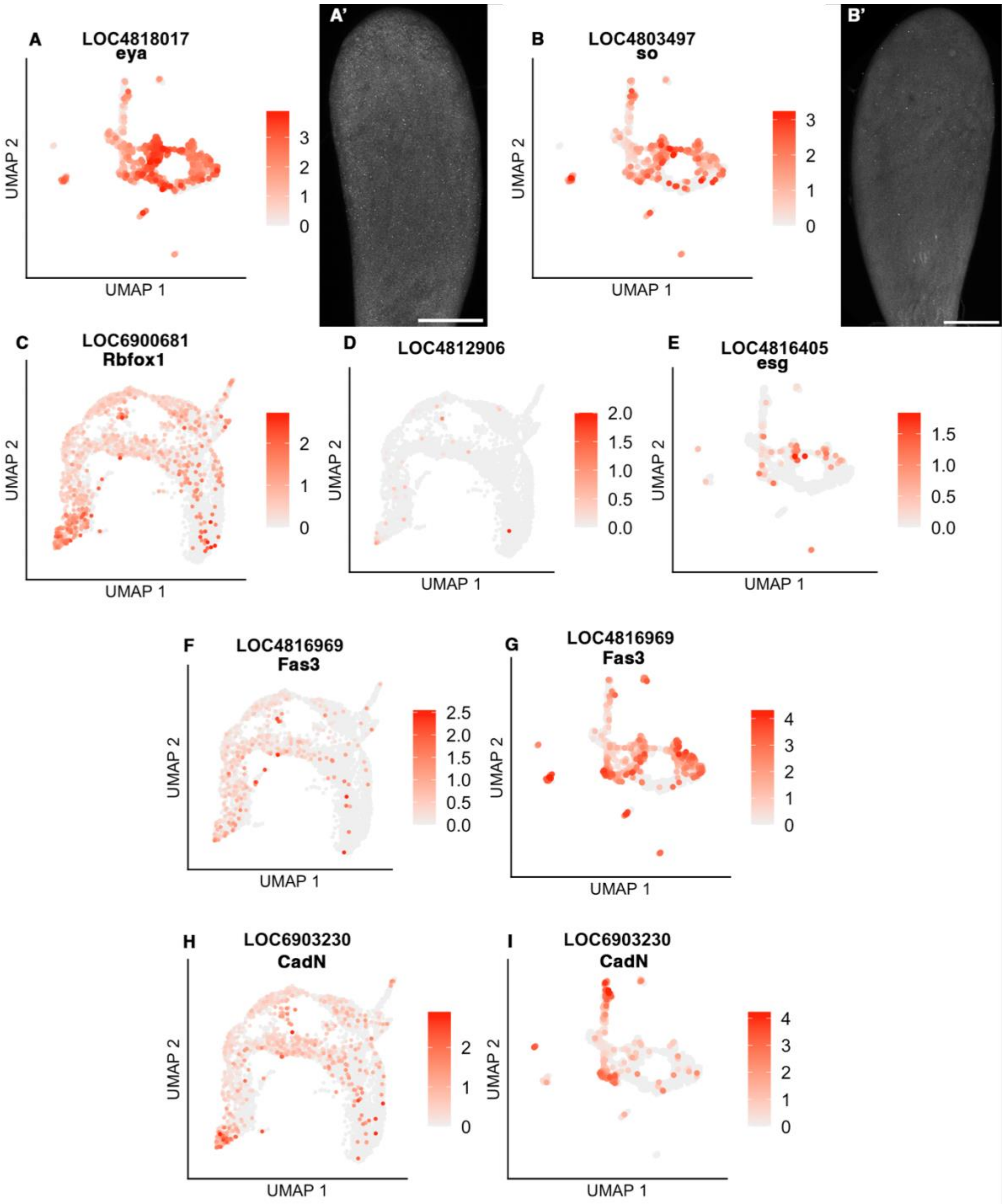


**Supplementary figure 8: Expression UMAPs of cyst cell marker genes.** (A) Somatic expression UMAP of *eyes absent (eya)* shows broad *eya* expression in somatic cell types. (A') HCR-FISH of *eya* showing broad low-level expression in somatic cells. (B) Somatic expression UMAP of *sine oculis (so)* shows similar broad cell-type expression to *eya*. (B') HCR-FISH of *so* shows low level expression, but cell types are not identifiable. (C) Germline expression UMAP of *Rbfox1* shows upregulation in the germline stem cells and spermatogonia. (D) Germline expression UMAP of *LOC4812906* does not indicate strong germline expression. (E) Somatic expression UMAP of *escargot (esg)* does not indicate upregulation in any one somatic cell type. (F-G) Germline and somatic expression UMAPs of *Fasciclin 3 (Fas3)* and (H-I) *N-Cadherin (CadN)*, which indicate the hub, cyst stem cells, and germline stem cells. *CadN* is also upregulated in muscle.

**Figure S9**

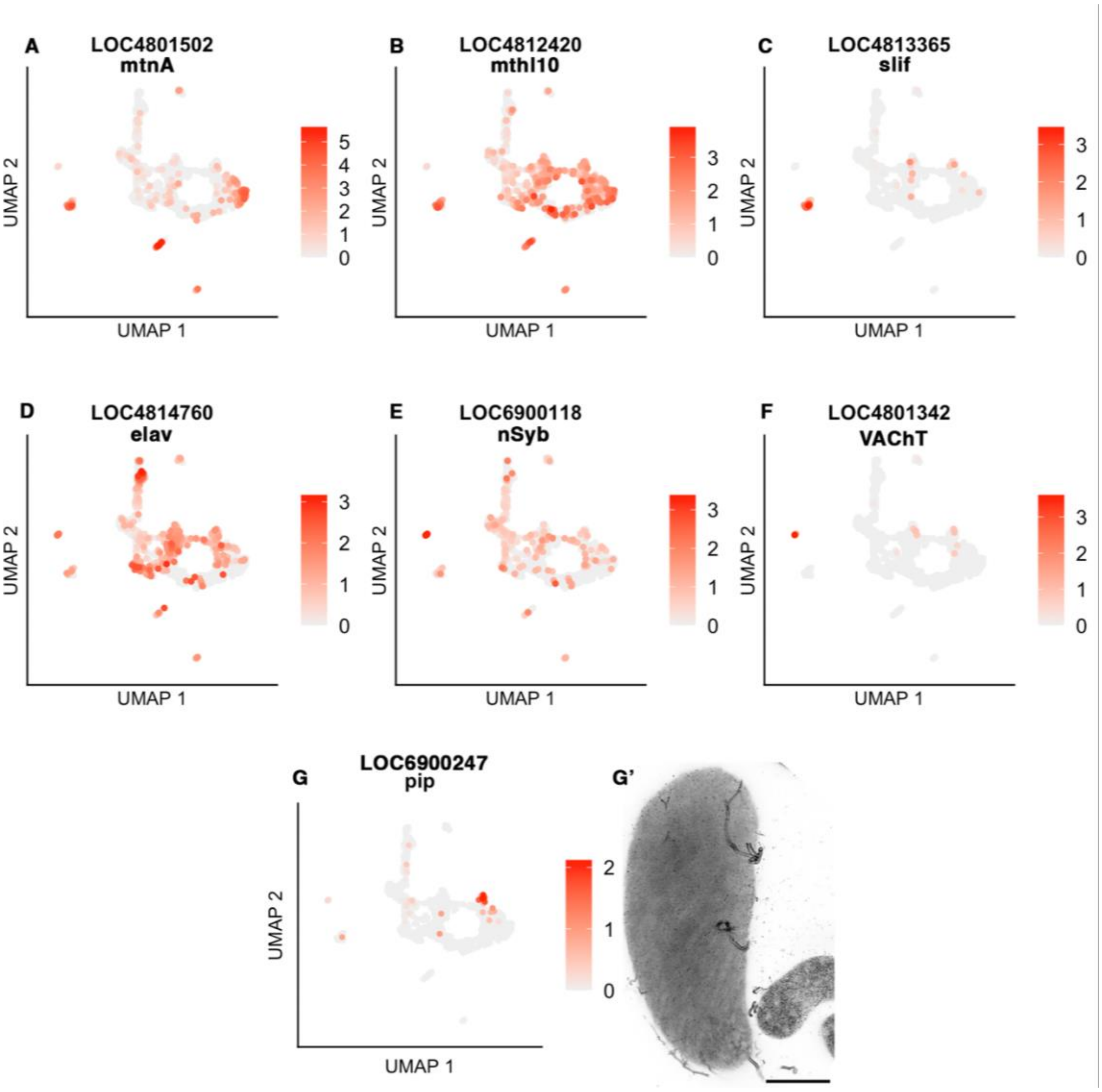


**Supplementary figure 9: Somatic expression UMAPs of non-cyst cell somatic cell marker genes.** (A-C) *Metallothionein A (MtnA), methuselah-like 10 (mthl10)* and *slimfast (slif)* are enriched in the terminal epithelium. (D-F) *elav, nSyb,* and *VAChT* are neuronal markers. (G, G') *pipe (pip)* is enriched in the testis sheath.
