## Supplementary material for "Single cell RNA sequencing of *D. pseudoobscura* testes reveals transcriptional signatures of heteromorphic spermatogenesis": Supplementary methods.docx

### Probe sequences for HCR-FISH validation of scRNAseq data and annotations.

| **Probe** | ***D. pse* Gene ID** | **Probe Sequences** | **Amplifier extension** | **Amplifier fluorophore** |
| --- | --- | --- | --- | --- |
| *esg* | LOC4816405 | GAGGAGGGCAGCAAACGGAAAGGCAAAGATCCTGGGGCTCATTGA | B1 |  |
|  |  | CAGCTCCTCCGACTCTGTTTTCTTATAGAAGAGTCTTCCTTTACG |  |  |
|  |  | GAGGAGGGCAGCAAACGGAAGCTGTAGGCCATACTTGGGATACGT |  |  |
|  |  | TACGGGGACATGTGGAAGTTCCATGTAGAAGAGTCTTCCTTTACG |  |  |
| *vasa* | LOC4817921 | CCTCGTAAATCCTCATCAAACACATTTGCCGCGATAATATCCCTC | B2 |  |
|  |  | TGGCCAACTTATAGCCCGATTTCGTAAATCATCCAGTAAACCGCC |  |  |
|  |  | CCTCGTAAATCCTCATCAAAAGATCTGAATGGCCAACTCGCGAGT |  |  |
|  |  | TGACTGAACTTCCTAGCCTCGTGATAAATCATCCAGTAAACCGCC |  |  |
|  |  | CCTCGTAAATCCTCATCAAATGGTGCCATCGGCGCTTTCTTTCAA |  |  |
|  |  | CCACGTTTGGTCTCAACGAAGACAAAAATCATCCAGTAAACCGCC |  |  |
| *aly* | LOC6900097 | GAGGAGGGCAGCAAACGGAATGGAAATTCTTGTGGCCGTGCCCAA | B1 |  |
|  |  | GCCTTTTCCTTTTCCACCTCCATCTTAGAAGAGTCTTCCTTTACG |  |  |
|  |  | GAGGAGGGCAGCAAACGGAAAAAGCGGGCCGTTTCACAAAGAGCT |  |  |
|  |  | GAACTTCGTTGGACTGAGCCCATAGTAGAAGAGTCTTCCTTTACG |  |  |
|  |  | GAGGAGGGCAGCAAACGGAAGCATATTCACCATTGTGCTGGCCAT |  |  |
|  |  | TTGAAAACGGCACTCAAATCGGGCGTAGAAGAGTCTTCCTTTACG |  |  |
|  |  | GAGGAGGGCAGCAAACGGAATCTTGCCTTCAAGGTAGGCTCCTTG |  |  |
|  |  | TTGGCTTTGGCTGTCTAGCAATTGGTAGAAGAGTCTTCCTTTACG |  |  |
| *protA* | LOC6903471 | GTCCCTGCCTCTATATCTTTCATTGCGCCTGAACTTTGATCTCTG | B3 |  |
|  |  | CCACTCCCATGGCAATCCATATCAATTCCACTCAACTTTAACCCG |  |  |
|  |  | GTCCCTGCCTCTATATCTTTTCCCCTCTTCCTCTTTCTTGAGTCT |  |  |
|  |  | CATTATTGGTAATGGGGCCAGGGTTTTCCACTCAACTTTAACCCG |  |  |
| *f-cup* | LOC6897136 | GAGGAGGGCAGCAAACGGAAGCCGTTGGATGGCGAAGTATTACAT | B1 |  |
|  |  | AGGAGCAGTCGTCAAAATCGTAGCCTAGAAGAGTCTTCCTTTACG |  |  |
|  |  | GAGGAGGGCAGCAAACGGAATTATACTGCTGCTCCATGGGTGGCT |  |  |
|  |  | GAGACCGGTGTACATCGATTGATGATAGAAGAGTCTTCCTTTACG |  |  |
|  |  | GAGGAGGGCAGCAAACGGAACAGATGCGGCAAGTTCTCACACATA |  |  |
|  |  | TATTGTCCCGGAGGTCGAGTACCTTTAGAAGAGTCTTCCTTTACG |  |  |
|  |  | GAGGAGGGCAGCAAACGGAAGCTGCTGTTGTTGATACATGGCTGA |  |  |
|  |  | ACGACGCACTGTCCGTAGAATAGTTTAGAAGAGTCTTCCTTTACG |  |  |
| *p-cup* | LOC4815368 | CCTCGTAAATCCTCATCAAAACTTCCTGACGATCCCATACTGCTC | B2 |  |
|  |  | TTGCGGGCGTGCATCATTTTCATGAAAATCATCCAGTAAACCGCC |  |  |
|  |  | CCTCGTAAATCCTCATCAAACCATCAGCTTGGTGGACTTTAGTTC |  |  |
|  |  | ATTGTGTTGCAGATGGCCAGGTAGGAAATCATCCAGTAAACCGCC |  |  |
| *caf1A* | LOC4803088 | GTCCCTGCCTCTATATCTTTTCCTCGACGGCATCGTCGAAAGATT | B3 |  |
|  |  | CTTGTACTCTTCGTTGATCACACGCTTCCACTCAACTTTAACCCG |  |  |
|  |  | GTCCCTGCCTCTATATCTTTAGCCATCAAACTGGGCATCTTCGCT |  |  |
|  |  | TCGCCTTTTTCATTGTCGTAGTGCGTTCCACTCAACTTTAACCCG |  |  |
|  |  | GTCCCTGCCTCTATATCTTTAATCCCAAACGTGCAAGCGACGATC |  |  |
|  |  | CTCTGCTCCTCGCCTATTTTCGACATTCCACTCAACTTTAACCCG |  |  |
|  |  | GTCCCTGCCTCTATATCTTTAAATGATCCAGGGCTCGTTGGGGTT |  |  |
|  |  | ATTATGTTGTCCTCGGACACGGAGCTTCCACTCAACTTTAACCCG |  |  |
| *caf1B* | LOC6897547 | CCTCAACCTACCTCCAACAATCTATCAATCTGTGCCTCCTCGGTG | B4 |  |
|  |  | CGCCCATCTCAATATCGTAACGCGAATTCTCACCATATTCGCTTC |  |  |
|  |  | CCTCAACCTACCTCCAACAAAGAGTCTTGTGGCATGTAGCGAGCA |  |  |
|  |  | TTGGCGATTTGGTGGCTATGATGCAATTCTCACCATATTCGCTTC |  |  |
|  |  | CCTCAACCTACCTCCAACAACGTGGTGTGCCTTAAGCGAATGTAA |  |  |
|  |  | TTCCAATGAACCTGGGTGATCTCTCATTCTCACCATATTCGCTTC |  |  |
|  |  | CCTCAACCTACCTCCAACAAATTGTCGGCGGACACAGAGCATATG |  |  |
|  |  | CGGCCATTTGCCACACTTCCATCAAATTCTCACCATATTCGCTTC |  |  |
| *comr* | LOC4805252 | CCTCGTAAATCCTCATCAAATTACCTCATCGTCGGCGATAGTTGC | B2 |  |
|  |  | ATCTTCTGGCGGACTCTAGTGAGGAAAATCATCCAGTAAACCGCC |  |  |
|  |  | CCTCGTAAATCCTCATCAAACCTTCGTCGACCAATTGAGGAAGTC |  |  |
|  |  | ACATACTCGAGCAGCAGGTGTTTGCAAATCATCCAGTAAACCGCC |  |  |
|  |  | CCTCGTAAATCCTCATCAAAACTATCGAGCTCTTGGGCAGATCCT |  |  |
|  |  | GAAGTTGGCATGCTGGAAGACCAAGAAATCATCCAGTAAACCGCC |  |  |
|  |  | CCTCGTAAATCCTCATCAAACGGATCTTTTTCGGCGCACCCATTA |  |  |
|  |  | TTGCGGCTTGAGGTTTCTTATGTGGAAATCATCCAGTAAACCGCC |  |  |
| *mip40* | LOC4804612 | GAGGAGGGCAGCAAACGGAACTTGTGCATCTTCTTGGGCGACTTC | B1 |  |
|  |  | TTAGCGAAAACTCAGCCCCCACACTTAGAAGAGTCTTCCTTTACG |  |  |
|  |  | GAGGAGGGCAGCAAACGGAAGCTCAAACAGGCGCATCACATACGA |  |  |
|  |  | TTGTACTTCGAAAGGTCCAGGCTGCTAGAAGAGTCTTCCTTTACG |  |  |
|  |  | GAGGAGGGCAGCAAACGGAATAATCTCGCCGCTCTTTAGCTTCGC |  |  |
|  |  | GGCTTCGGCATGTGGGTCAGAATTTTAGAAGAGTCTTCCTTTACG |  |  |
|  |  | GAGGAGGGCAGCAAACGGAATATGCTCTTCCCAGTGATCGCGTAC |  |  |
|  |  | TAGCGCTCGTGGTTGTACTTTTGAGTAGAAGAGTCTTCCTTTACG |  |  |
| *TGIF (achi/vis)* | LOC4805467 | CCTCGTAAATCCTCATCAAATGTAGACAACGCTCTCGTCGTACTC | B2 |  |
|  |  | GATTGTTTGCAACGGCCTGCTGCCAAAATCATCCAGTAAACCGCC |  |  |
|  |  | CCTCGTAAATCCTCATCAAAGTGCTGGTAAAGGGCTTTATGGCAC |  |  |
|  |  | AAACTCTGCCGGCATCTGCCGAAATAAATCATCCAGTAAACCGCC |  |  |
|  |  | CCTCGTAAATCCTCATCAAATTTGTTAGCCACCTCCGAGCGCTTA |  |  |
|  |  | CTGCTTTTTGCTCGGATTGAAGCGTAAATCATCCAGTAAACCGCC |  |  |
| *tomb* | LOC4818077 | GTCCCTGCCTCTATATCTTTCACATCCCTTGATGGTGGACTCCTT | B3 |  |
|  |  | TTAATGCACGAGGTACGCTTGCACGTTCCACTCAACTTTAACCCG |  |  |
|  |  | GTCCCTGCCTCTATATCTTTCGCTCCACCGAGTTTTTGCAATCAA |  |  |
|  |  | TGGCAGATTGTCGGTAACAAGCGGATTCCACTCAACTTTAACCCG |  |  |
|  |  | GTCCCTGCCTCTATATCTTTTCTTGGGGCATTCCACGTGGTTTGT |  |  |
|  |  | CTGTGGCATTTGGATTATAACCGGCTTCCACTCAACTTTAACCCG |  |  |
|  |  | GTCCCTGCCTCTATATCTTTGGGCTGTATGAACAGATTGAGTTCC |  |  |
|  |  | TGCACTCGAGCAGTGTCGTATTGACTTCCACTCAACTTTAACCCG |  |  |
| *topi* | LOC4801505 | CCTCAACCTACCTCCAACAACTCTGTGGTCATCTTACTGTTGGTG | B4 |  |
|  |  | TGCGTTGTGCTTGAAAGACCGACTCATTCTCACCATATTCGCTTC |  |  |
|  |  | CCTCAACCTACCTCCAACAAATGATGCGCTCGGTGCTTCGAACAT |  |  |
|  |  | GCGTAGGCGTACTTGATTTTCCAAGATTCTCACCATATTCGCTTC |  |  |
|  |  | CCTCAACCTACCTCCAACAAGGTCGTGTTCATTGTAATGCCGCAG |  |  |
|  |  | TCTGGAAATGGACACTGGCCTCCTTATTCTCACCATATTCGCTTC |  |  |
|  |  | CCTCAACCTACCTCCAACAAATCTTCCTCAGGACCGACGTGAGAT |  |  |
|  |  | CTTCCGTAGCGAAGTTGTTTGGACCATTCTCACCATATTCGCTTC |  |  |
| *wuc* | LOC4804758 | GTCCCTGCCTCTATATCTTTTTCAGCTATGATGGAGCTGACGCCA | B3 |  |
|  |  | CTTCTTCCTCGTCCTTCTCCTTCATTTCCACTCAACTTTAACCCG |  |  |
|  |  | GTCCCTGCCTCTATATCTTTCCTCAGTAAATCCTGCCTTGGCGTT |  |  |
|  |  | AGTTCCTCCATGGTCTTGACGTCCTTTCCACTCAACTTTAACCCG |  |  |
| *kmg* | LOC4816438 | GAGGAGGGCAGCAAACGGAAGTAATGACATGTTAGCCGGATCCGC | B1 |  |
|  |  | TGGTACCATGCGACTTCCTGTGAGATAGAAGAGTCTTCCTTTACG |  |  |
|  |  | GAGGAGGGCAGCAAACGGAAATGATGATCCTTTCGCCGTGGACGT |  |  |
|  |  | ATAGAGGCACAGCTTGCACTCGTTTTAGAAGAGTCTTCCTTTACG |  |  |
|  |  | GAGGAGGGCAGCAAACGGAAACCAGTTTCTGGAAATGACGCGAGG |  |  |
|  |  | GCCATGCACCATTTTCATGTGACGCTAGAAGAGTCTTCCTTTACG |  |  |
|  |  | GAGGAGGGCAGCAAACGGAAGCTGAAACTCAAAGGCCATCGAGAA |  |  |
|  |  | TCCATCATGGGTGGCATCATCTTGCTAGAAGAGTCTTCCTTTACG |  |  |
| *exd* | LOC4811784 | GAGGAGGGCAGCAAACGGAAAATCTGCGCCAACTTTGCGCGATAG | B1 |  |
|  |  | CTAGCTCTTGGTGATAGATCTGACGTAGAAGAGTCTTCCTTTACG |  |  |
|  |  | GAGGAGGGCAGCAAACGGAATGAACTCGTTACAAGCCTGCTCGTA |  |  |
|  |  | CGCAGAAGATTCATAACGTGCGTAGTAGAAGAGTCTTCCTTTACG |  |  |
|  |  | GAGGAGGGCAGCAAACGGAAATTACAGCTTCGCATGTCGACTGCT |  |  |
|  |  | GTCTAGAAAGCGCGAACGAAGAATCTAGAAGAGTCTTCCTTTACG |  |  |
| *hth* | LOC4801015 | GAGGAGGGCAGCAAACGGAATACATGACTAGGTGTACCATGACCC | B1 |  |
|  |  | CCATTAGGTGATTACCAACCGGTGATAGAAGAGTCTTCCTTTACG |  |  |
|  |  | GAGGAGGGCAGCAAACGGAAATATGAGCGCCAATAGCGGGAATAG |  |  |
|  |  | CATGTGGCCAATTCGCATTTCTCGATAGAAGAGTCTTCCTTTACG |  |  |
|  |  | GAGGAGGGCAGCAAACGGAAAGACTGAGGAATACAGTCGCCTGGA |  |  |
|  |  | CCAAAGTTGTCAGGACTACCAAGGATAGAAGAGTCTTCCTTTACG |  |  |
|  |  | GAGGAGGGCAGCAAACGGAAACAGCCACGCTCTCAATATGTTGGT |  |  |
|  |  | GGGTAGGGATGCGTTAAATGCTGAATAGAAGAGTCTTCCTTTACG |  |  |
| *br* | LOC6902077 | GAGGAGGGCAGCAAACGGAAACAATTCGCGGAAGTAGGGACTGCA | B1 |  |
|  |  | GGATGTTTGCAGGGTGTGCTCTTCATAGAAGAGTCTTCCTTTACG |  |  |
|  |  | GAGGAGGGCAGCAAACGGAATATGCGTATCCTCTGCCTGCTGTTG |  |  |
|  |  | AGGTTCTGTATTTGAGCCAGGTGGCTAGAAGAGTCTTCCTTTACG |  |  |
|  |  | GAGGAGGGCAGCAAACGGAATGGGCTCTGACTTGACCTGATCATC |  |  |
|  |  | TTGTTGTTGGAGCACACCAGCTCCATAGAAGAGTCTTCCTTTACG |  |  |
| *Ets98B* | LOC6896869 | GAGGAGGGCAGCAAACGGAATTGTACTGGTACACCGTGGTGGGTA | B1 |  |
|  |  | GCATTCAATCTTAATCTGTGGCGCGTAGAAGAGTCTTCCTTTACG |  |  |
|  |  | GAGGAGGGCAGCAAACGGAATACAGATGGCATCAGCTTCCCGCTT |  |  |
|  |  | GGATCCTGCGATATTTGCAGCTCCATAGAAGAGTCTTCCTTTACG |  |  |
|  |  | GAGGAGGGCAGCAAACGGAAAGGTTCAGCATCGGATAGCTAGCTG |  |  |
|  |  | TTCGCTGTCCTCATTGTAGTCGTCCTAGAAGAGTCTTCCTTTACG |  |  |
| *fkh* | LOC4801871 | GAGGAGGGCAGCAAACGGAATGATCAGGGAGATGTAGCTGTACGG | B1 |  |
|  |  | GTCGGATTATTCTGTATGGCCATCGTAGAAGAGTCTTCCTTTACG |  |  |
|  |  | GAGGAGGGCAGCAAACGGAAAAATCTCCGACAGAGTGAGCATCCG |  |  |
|  |  | GGGAACAGGTCCATGATGAACTGGTTAGAAGAGTCTTCCTTTACG |  |  |
|  |  | GAGGAGGGCAGCAAACGGAAGATGCAGTGTCCAGAAGGATCCCTT |  |  |
|  |  | TTCTCGAACATATTCCCGGAGTCGGTAGAAGAGTCTTCCTTTACG |  |  |
| *tj* | LOC4816276 | GAGGAGGGCAGCAAACGGAATTAACGGTGGCGAATGAGTCTGGCT | B1 |  |
|  |  | GCCCACTGCGTCCAATTTCAAAGAGTAGAAGAGTCTTCCTTTACG |  |  |
|  |  | GAGGAGGGCAGCAAACGGAACGTGTCGGGCCGAAGTTTTTCTTTA |  |  |
|  |  | GCACTTAGCACTCGGATACTCGTACTAGAAGAGTCTTCCTTTACG |  |  |
|  |  | GAGGAGGGCAGCAAACGGAAATTTTCATTGGATCGACCGCGACCG |  |  |
|  |  | CGTATCGGCTATAGTGGGATCTTCCTAGAAGAGTCTTCCTTTACG |  |  |
| *LOC4812906* | LOC4812906 | CCTCGTAAATCCTCATCAAACTGCTTGGCCTTCATTGTCTTCACG | B2 |  |
|  |  | AGAGGACAAACTCCACATTGTCGCCAAATCATCCAGTAAACCGCC |  |  |
|  |  | CCTCGTAAATCCTCATCAAACGGCACATCTTGATATCGAGCACCA |  |  |
|  |  | AAAGCCCCAGGGCACATTGTCAAAGAAATCATCCAGTAAACCGCC |  |  |
|  |  | CCTCGTAAATCCTCATCAAAGTTTCGTCTTGGTCACCTCCTCGTT |  |  |
|  |  | TTGCGGTTCTCCTTGTCCTTCTCCAAAATCATCCAGTAAACCGCC |  |  |
| *Rbfox1* | LOC6900681 | CCTCGTAAATCCTCATCAAAATTAAATGTGGCGGCTGCTACTGCG | B2 |  |
|  |  | ACATGCTGCTATCTGTGGCACTTCCAAATCATCCAGTAAACCGCC |  |  |
|  |  | CCTCGTAAATCCTCATCAAATGCTGTTGCTGCAGTTGTAGCTGGT |  |  |
|  |  | TTGTTGTTGTTGGTGGTGCTGCTGCAAATCATCCAGTAAACCGCC |  |  |
|  |  | CCTCGTAAATCCTCATCAAAGTGACCACTGAAACTGGGATTTGGC |  |  |
|  |  | ATGTCCATTTGTCAGGGATGGCTGCAAATCATCCAGTAAACCGCC |  |  |
|  |  | CCTCGTAAATCCTCATCAAACGATTTAATATGGCGGCTGGCTGAG |  |  |
|  |  | ACTGTTTAGCGTAGTCACAGTGCGGAAATCATCCAGTAAACCGCC |  |  |
| *LOC4803306* | LOC4803306 | GTCCCTGCCTCTATATCTTTAGACGAGTAGCAATGCTGTCAAGCA | B3 |  |
|  |  | CTCCCAAGTCCGTAAGGGAGAAAAATTCCACTCAACTTTAACCCG |  |  |
|  |  | GTCCCTGCCTCTATATCTTTGTATGAGCCTGGTATTGGGCATACA |  |  |
|  |  | AGGGATATGCGGTAGATTGTCATGGTTCCACTCAACTTTAACCCG |  |  |
| *Ggt-1* | LOC4814339 | CCTCGTAAATCCTCATCAAAACCAGTTCGGGGCTGAGGATTTTCT | B2 |  |
|  |  | ATGAATGAGCTTGCGTACATCGGCCAAATCATCCAGTAAACCGCC |  |  |
|  |  | CCTCGTAAATCCTCATCAAATTATGACCATTGCCACGCTTGTGGT |  |  |
|  |  | GTCTCGTGGCGAATTAGGTACTTCAAAATCATCCAGTAAACCGCC |  |  |
| *LOC117183237* | LOC117183237 | GTCCCTGCCTCTATATCTTTACACTCCATCAAAAAAGCGCTGTGG | B3 |  |
|  |  | AAACTCGACCTCTGTTGTGGCATCATTCCACTCAACTTTAACCCG |  |  |
|  |  | GTCCCTGCCTCTATATCTTTCCACAGCTTTGAAGTGTCGTTTCGT |  |  |
|  |  | TTTAGCGTAGAAGCCCCAGCCACATTTCCACTCAACTTTAACCCG |  |  |
| *Idgf4* | LOC4814222 | GAGGAGGGCAGCAAACGGAAGATCCAGATCGTTGAGGGTGAGCTT | B1 |  |
|  |  | AAATGGGTGCAGTACTGCAAGGCAGTAGAAGAGTCTTCCTTTACG |  |  |
|  |  | GAGGAGGGCAGCAAACGGAACATCGAAGAAAAGGGAGGAGTTGAC |  |  |
|  |  | TCCAAATTGTTGATCACGGCCGGGATAGAAGAGTCTTCCTTTACG |  |  |
|  |  | GAGGAGGGCAGCAAACGGAATAGTTGACATTCAGCTCGGGGTTGC |  |  |
|  |  | GTTGTTCACCCAGTACTTGACCTGGTAGAAGAGTCTTCCTTTACG |  |  |
| *arqs* | LOC6896834 | GAGGAGGGCAGCAAACGGAACCAGTAGCAGCAACAGGTAAATGCT | B1 |  |
|  |  | GCATTGATTTGCAGGCATTGCAGGCTAGAAGAGTCTTCCTTTACG |  |  |
|  |  | GAGGAGGGCAGCAAACGGAATCCTTATCGAGCTTATCGCGCGCAT |  |  |
|  |  | AAATTCGACTGTCCTCTCTACTGCCTAGAAGAGTCTTCCTTTACG |  |  |
|  |  | GAGGAGGGCAGCAAACGGAACGATGTCGTTTGAGGTGCCGTTTTC |  |  |
|  |  | AACTTCTTCGCCAGGGAGATCAAGGTAGAAGAGTCTTCCTTTACG |  |  |
|  |  | GAGGAGGGCAGCAAACGGAAGTGCTTCAGATTGCTCACACAGTAG |  |  |
|  |  | GGGTGCTGCAGGCCGTATTTAGATTTAGAAGAGTCTTCCTTTACG |  |  |
| *pip* | LOC6900247 | GAGGAGGGCAGCAAACGGAAACGGTATGCGGCGATATTTGAAGGC | B1 |  |
|  |  | ATCAGTTCCACGGATCGCTTCGGATTAGAAGAGTCTTCCTTTACG |  |  |
|  |  | GAGGAGGGCAGCAAACGGAAATATCGTCGTGTTTGCCCGCATCGT |  |  |
|  |  | TTGGTAGGTGCCATGCAGAGTGTTGTAGAAGAGTCTTCCTTTACG |  |  |

### Primer sequences for piggyBac-Kmg-GFP construct assembly

| **Fragment** | **Forward primer** | **Reverse primer** |
| --- | --- | --- |
| *kmg* Promotor, 5’ UTR, CDS | CCTAGGCTTTTGCAGGAGGTTTCGCC | CCTAGGATATTGTTCATCACCGGCGGG |
| *kmg* 3’ UTR | GCGGCCGCTTAGAACCGGACTCTGACTGAAAAGGG | GCGGCCGCGCCGCTAGTCGTCTTAATATG |
| GFP C-tag | CCTAGGAGTAAAGGAGAAGAACTTTTCACTGG | GCTCTCCCATATGGTCGACCTG (NotI restriction site from pGEM-T Easy) |

### piggyBac-Kmg-GFP Construct Map

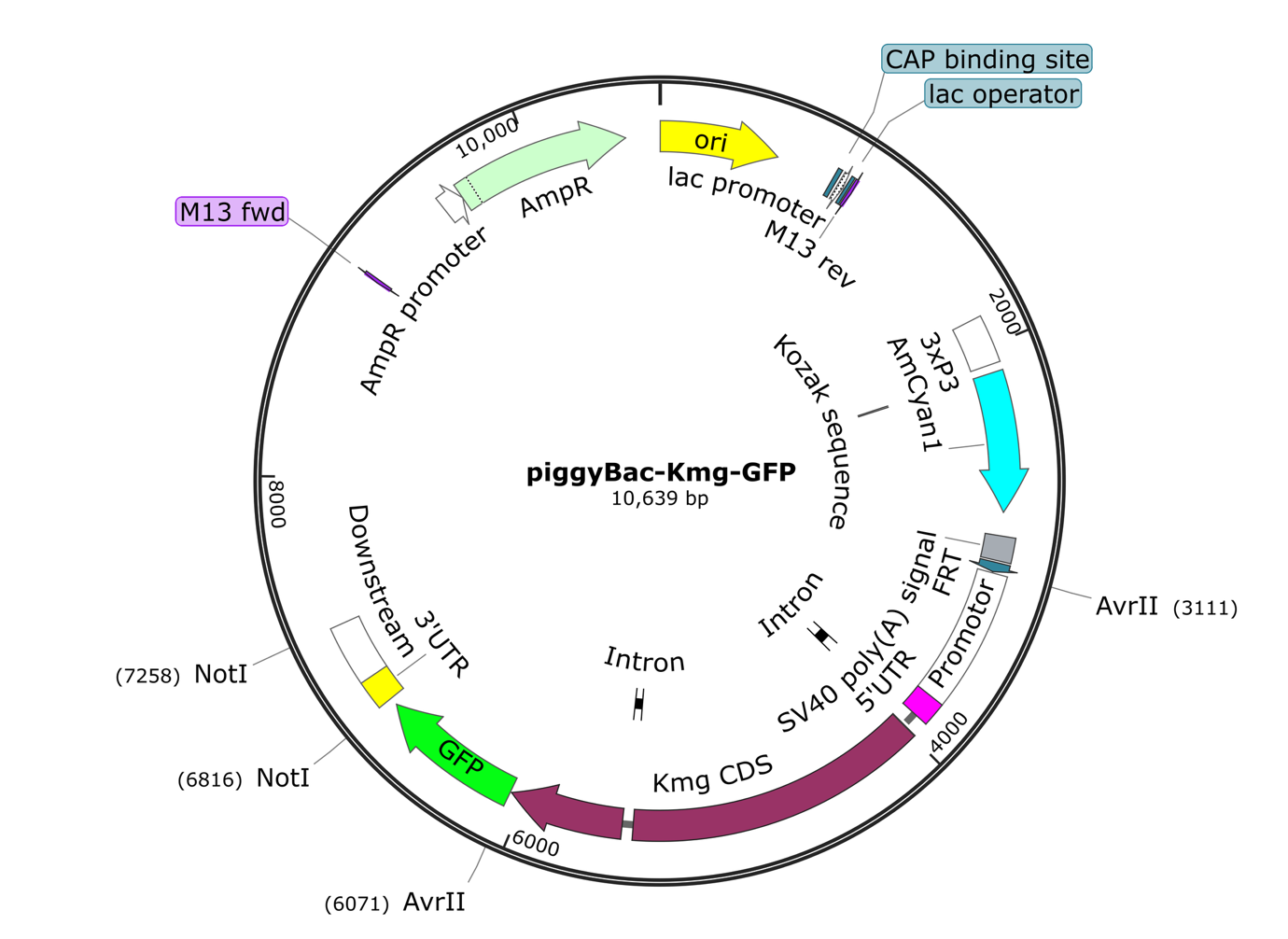
