## Supplementary material for "Single cell RNA sequencing of *D. pseudoobscura* testes reveals transcriptional signatures of heteromorphic spermatogenesis": Supplementary methods.pdf

### Primer sequences for piggyBac-Kmg-GFP construct assembly

| Fragment | Forward primer | Reverse primer |
|---|---|---|
| *kmg* Promotor, 5' UTR, CDS | CCTAGGCTTTTGCAGGAG GTTTCGCC | CCTAGGATATTGTTCATCAC CGGCGGG |
| *kmg* 3' UTR | GCGGCCGCTTAGAACCG GACTCTGACTGAAAAGGG | GCGGCCGCGCCGCTAGT CGTCTTAATATG |
| GFP C-tag | CCTAGGAGTAAAGGAGAA GAACTTTTCACTGG | GCTCTCCCATATGGTCGAC CTG (NotI restriction site from pGEM-T Easy) |

### piggyBac-Kmg-GFP Construct Map

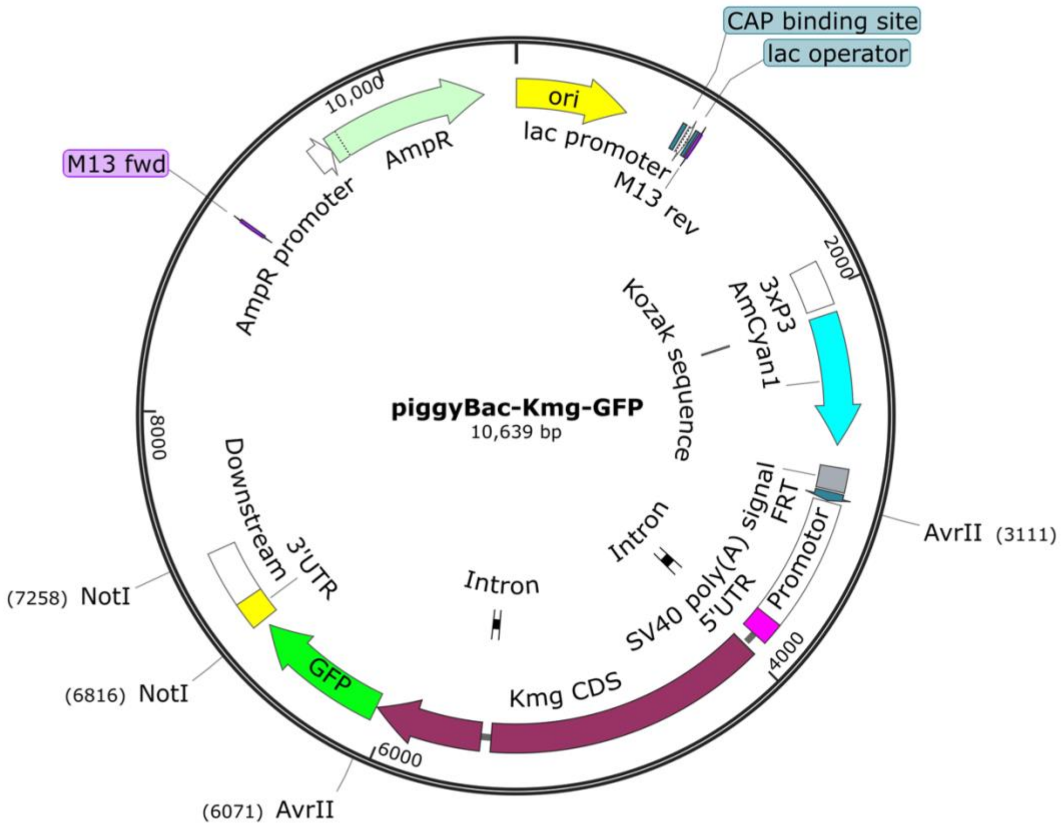
