## Supplementary material for "Single cell RNA sequencing of *D. pseudoobscura* testes reveals transcriptional signatures of heteromorphic spermatogenesis": Table S2 + S3 - Cell counts.pdf

**Supplementary table 2: Germline cluster transcript counts, feature counts, and number of cells per replicate, at clustering resolution Louvain 0.8.** Total cell counts per replicate are given.

| Germline Cluster (Louv 0.8) | Mean | | Median | | Cells per Replicate | |
|---|---|---|---|---|---|---|
| | Transcript Count | Feature Count | Transcript Count | Feature Count | Rep 1 | Rep 2 |
| **0** | 803.89 | 561.78 | 668 | 501 | 353 | 503 |
| **1** | 1421.39 | 846.83 | 960.5 | 703.5 | 274 | 404 |
| **2** | 2686.24 | 1159.74 | 2188.5 | 1094 | 280 | 368 |
| **3** | 8464.85 | 2340.53 | 6464 | 2267 | 238 | 235 |
| **4** | 18640.04 | 3365.16 | 12928.5 | 3205.5 | 204 | 240 |
| **5** | 51988.41 | 5765.4 | 37101 | 5837 | 126 | 317 |
| **6** | 3494.61 | 1311.16 | 2218.5 | 1101.5 | 203 | 225 |
| **7** | 25752 | 4445.35 | 23765 | 4508.5 | 196 | 182 |
| **8** | 12417.12 | 3235.65 | 8380.5 | 3089 | 112 | 230 |
| **9** | 58077.36 | 5591.92 | 53567 | 6023 | 207 | 120 |
| **10** | 7996.08 | 1675.78 | 4077.5 | 1412 | 121 | 109 |
| **11** | 24082.88 | 4050.93 | 11038 | 3653 | 75 | 64 |
| | | | | **Total** | **2389** | **2997** |

**Supplementary table 3: Somatic cluster transcript counts, feature counts, and number of cells per replicate, at clustering resolution Louvain 0.8.** Total cell counts per replicate are given.

| Somatic Cluster (Louv 0.8) | Mean | | Median | | Cells per Replicate | |
|---|---|---|---|---|---|---|
| | Transcript Count | Feature Count | Transcript Count | Feature Count | Rep 1 | Rep 2 |
| **0** | 7204.89 | 2304.74 | 6225 | 2145 | 64 | 115 |
| **1** | 4257.93 | 1506.30 | 2134 | 1209.5 | 18 | 126 |
| **2** | 3313.38 | 1245.09 | 1464 | 874.5 | 43 | 83 |
| **3** | 19065.01 | 3018.82 | 5152.5 | 2005.5 | 23 | 99 |
| **4** | 58635.36 | 6125.23 | 36671 | 5914 | 14 | 97 |
| **5** | 8507.29 | 2443.15 | 4638.5 | 1995.5 | 42 | 68 |
| **6** | 10855.77 | 2825.16 | 9797.5 | 2700.5 | 47 | 47 |
| **7** | 12120.84 | 3005.53 | 9043.5 | 2738.5 | 7 | 73 |
| **8** | 1822.87 | 973.53 | 891.5 | 635 | 26 | 50 |
| **9** | 16497.88 | 2816.06 | 5204.5 | 2193.5 | 19 | 45 |
| **10** | 11194.24 | 2477.34 | 6594 | 2237 | 12 | 17 |
| **11** | 17136.23 | 3039.69 | 11519 | 2708 | 10 | 3 |
| | | | | **Total** | **325** | **823** |
