## Supplementary material for "Single cell RNA sequencing of *D. pseudoobscura* testes reveals transcriptional signatures of heteromorphic spermatogenesis": Table S4 - Cluster annotations.pdf

**Supplementary Table 1: Germline cluster annotations at Louvain resolutions 0.8, 1.6 and 3.2.** Cell type annotations of clusters and morph are described. **GSC** Germline stem cell. **SG** spermatogonia. **SC** spermatocyte. **EuSC** euspermatocyte. **EuST** euspermatid. **PaSC** paraspermatocyte. **PaST** paraspermatid.

| Louvain Resolution | Cluster | Annotation | Broad Annotation | Morph |
|---|---|---|---|---|
| 0.8 | 11 | GSC-SG | GSC-SG | Pre-trajectory split |
| | 8 | SG-SC | SG-SC | |
| | 5 | SC (Early) | SC | |
| | 9 | EuSC (Mid) | EuSC | Eusperm |
| | 4 | EuSC (Late) | | |
| | 6 | EuST (Early) | EuST | |
| | 7 | PaSC (Mid) | PaSC | Parasperm |
| | 3 | PaSC (Late) | | |
| | 2 | PaST (Early) | PaST | |
| | 10 | Spermatid | Spermatid | Both |
| 1.6 | 7 | GSC-SG | GSC-SG | Pre-trajectory split |
| | 12 | SG-SC | SG-SC | |
| | 13 | SC (Early) | SC | |
| | 15 | EuSC (Early) | EuSC | Eusperm |
| | 8 | EuSC (Mid) | | |
| | 5 | EuSC (Late) | | |
| | 4 | EuST (Early) | EuST | |
| | 11 | EuST (Mid) | | |
| | 17 | EuST (Late) | | |
| | 9 | PaSC (Early) | PaSC | Parasperm |
| | 10 | PaSC (Mid) | | |
| | 2 | PaSC (Late) | | |
| | 3 | PaST (Early) | PaST | |
| | 6 | PaST (Mid) | | |
| | 14 | PaST (Late) | | |
| 3.2 | 18 | GSC-SG | GSC-SG | Pre-trajectory split |
| | 16 | SG | SG | |
| | 11 | SC (Early) 1 | SC | |
| | 13 | SC (Early) 2 | | |
| | 21 | EuSC (Early) | EuSC | eusperm |
| | 19 | EuSC (Mid) 1 | | |
| | 15 | EuSC (Mid) 2 | | |

| | | | | |
|---|---|---|---|---|
| | 27 | EuSC (Late) 1 | | |
| | 9 | EuSC (Late) 2 | | |
| | 22 | EuSC Meiotic | | |
| | 7 | EuST (Early) 1 | EuST | |
| | 6 | EuST (Early) 2 | | |
| | 30 | EuST (Mid) | | |
| | 26 | EuST (Late) | | |
| | 20 | PaSC (Early) | PaSC | Parasperm |
| | 17 | PaSC (Mid) 1 | | |
| | 14 | PaSC (Mid) 2.1 | | |
| | 25 | PaSC (Mid) 2.2 | | |
| | 10 | PaSC (Late) 1 | | |
| | 5 | PaSC Meiotic | | |
| | 8 | PaST (Early) 1 | PaST | |
| | 4 | PaST (Early) 2 | | |
| | 12 | PaST (Early) 3 | | |
| | 23 | PaST (Late) | | |
